# Metabolic, epigenetic and transcriptomic alterations in postnatal 16p11.2-deficient murine astrocytes

**DOI:** 10.64898/2025.12.16.694710

**Authors:** Nicole Blakeley, Shama Naz, Ziying Liu, Tyler Renner, Sonia Leclerc, Cesar H. Comin, Matheus V. da Silva, Caroline Vergette, Christopher J. Porter, Theodore J. Perkins, Baptiste Lacoste

## Abstract

Autism Spectrum Disorders (ASD) are associated with metabolic dysregulation. While astrocytes are integral to cerebral metabolism, their molecular and functional changes in ASD are poorly known. Using early postnatal primary cortical astrocytes from a mouse model of 16p11.2 deletion ASD syndrome (*16p11.2^df/+^* mice), we observed core molecular alterations with sex-specific profiles, suggesting divergent energetic pathways and epigenetic regulation. Targeted metabolomics revealed opposing phenotypes in male versus female *16p11.2^df/+^* astrocytes, particularly for alpha-ketoglutaric acid. Functionally, *16p11.2^df/+^*astrocytes exhibited elevated phosphorylation in low glucose culture conditions, and reduced glycolysis in high glucose. Epigenetic profiling of male *16p11.2^df/+^*astrocytes revealed differentially hydroxymethylated and methylated regions, with foci on chromosomes 3 and 13. Finally, bulk RNA sequencing in male and female mutant astrocytes indicated differential gene expression with profound sex differences, mostly affecting pathways related to cellular morphology. By establishing *16p11.2^df/+^* astroglial molecular signatures, this study refines our understanding of glial changes in ASD.

## INTRODUCTION

Accounting for 20 to 40% of brain cells and responsible for many roles^1^, astrocytes are integral to maintaining cerebral health. A principal function of astrocytes is forming communicative networks between various cell types including neurons and vascular endothelial cells (ECs). For instance, astrocyte sensing of increased neuronal activation via glutamate signaling induces ECs to modulate cerebral blood flow^2^. Astrocytes are a center piece in the cerebral energetics machinery, providing fuels to neurons^3–6^, and they are vulnerable to injury, inflammation and metabolic shortfalls, altering disease onset and/or pathogenesis^7^. Although limited research has focused on astrocytes in Autism Spectrum Disorders (ASD), mounting evidence suggests changes in astrocytes function, glial metabolism, and overall brain metabolism^8^. The link between astrocytes and ASD is mostly supported by changes to calcium signaling, process that may impact mitochondrial function^9^, gene expression^10^, and consequently neuronal function^10,11^. Disrupted calcium signaling was connected to morphological changes in astrocytes resultant from cytoskeletal alterations^12^. In a mouse model of the 16p11.2 deletion ASD Syndrome (*16p11.2^df/+^* in mice), we reported an increase in brain glucose uptake using F-Fluorodeoxyglucose positron emission tomography and intracerebral glucose sensor electrodes, as well as shift in brain glucose metabolism via metabolomics^13^. Other astrocyte specific metabolic dysfunctions linked to ASD include transporter and enzyme alterations. A reduction of glutamate transporter 1 (GLT-1) in astrocytes paired with glutamate/GABA imbalance was associated with neuronal apoptosis in rodent ASD models^14–16^. Astrocytic glutamate imbalance led to pathological repetitive behaviour in ASD mouse models such as excessive grooming^16^. Furthermore, disruptions in astrocyte calcium signaling failed to initiate astrocytic release of ATP in mice, leading to ASD behavioural phenotypes^10^. Knocking out type 2 inositol 1,4,5-trisphosphate receptor (IP3R2) in a mouse model disrupted astrocytic calcium signaling, leading to repetitive behaviours and social deficits. Additionally, the implantation in healthy mice of ASD patient-derived astrocytes exhibiting altered calcium signaling resulted in impaired memory and repetitive behaviour^17^. Such findings suggest that metabolic dysfunctions and signaling disruptions in astrocytes are involved in ASD pathophysiology. However, how astroglial molecular signatures are altered in critical developmental stages in ASD has yet to be elucidated. To address this gap in knowledge, we have undertaken a detailed profiling of primary murine cortical astrocytes extracted from male and female *16p11.2^df/+^*and WT littermates at postnatal day (P)8, at a time when astrocytes enter active morphogenesis including endfoot placement around the vasculature^18^. We show that in early postnatal development, *16p11.2^df/+^* astrocytes exhibit metabolic, epigenetic and transcriptomic alterations with pronounced sex-differences, altogether furthering our knowledge on glial changes in ASD.

## RESULTS

### Sex-specific differences in intracellular metabolite abundance between WT and *16p11.2^df/+^* astrocytes

We first aimed to assess the impact of 16p11.2 (7qF3) haploinsufficiency (Figure S1A) on intracellular metabolite abundance in cultured primary cortical astrocytes isolated from male and female WT and *16p11.2^df/+^* littermates at P8, using a targeted liquid chromatography-mass spectrometry (LC-MS) approach (Figure S1B). LC-MS detected a total of 104 metabolites. Overall, male *16p11.2^df/+^* astrocytes displayed an increase in metabolite concentrations compared to WT cells (Figure 1A) except for adenosine. Female *16p11.2^df/+^*astrocytes displayed a distinct profile, with predominantly reduced metabolite abundance (Figure 1B) with only a cluster of increased metabolites. These sex-specific differences were driven by a subset of metabolites with higher variable importance plot (VIP) scores (Figure 1C,D). Strong separation between WT and *16p11.2^df/+^* samples was confirmed by partial least squares discriminant analyses (PLS-DA) (Figure 1E,F).

**Figure 1.**
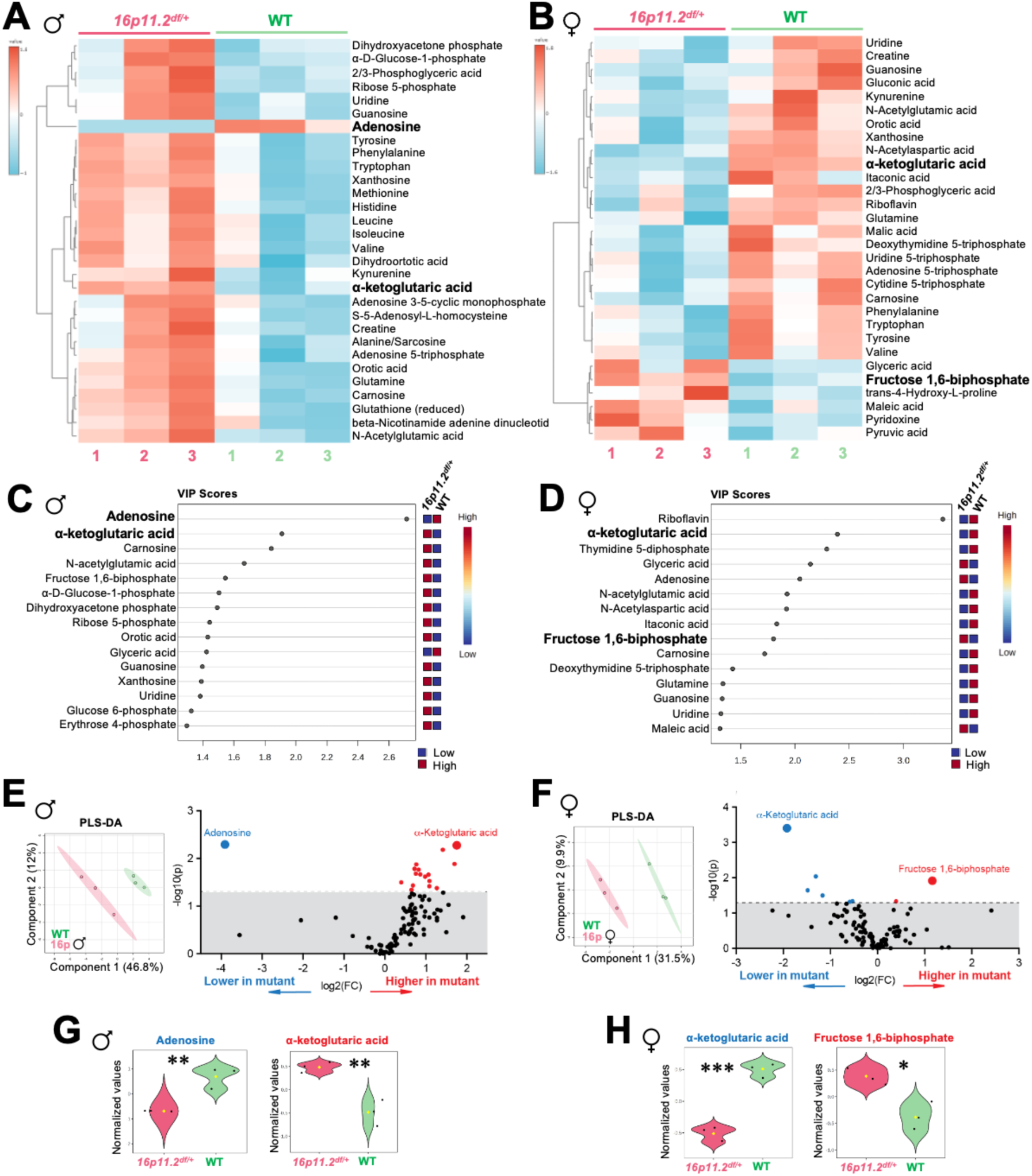
Sex-specific metabolomic profiles in cortical astrocytes from 16p11.2-deficient versus WT P8 pups. (A) Heat map (multivariate statistical analysis) displaying the top 30 metabolites significantly altered by genotype from male *16p11.2^df/+^* and WT cortical astrocytes, isolated from 3 mice (n=3) per genotype. (B) Heat map (multivariate statistical analysis) displaying the top 30 metabolites significantly altered by genotype from female *16p11.2^df/+^* (n=3) and WT (n=3) cortical astrocytes. (C) Variable importance in projection (VIP) scores, showing the top 15 (score > 1) metabolite drivers contributing to the separation of metabolic profiles identified by Partial Least Squares-Discriminant Analysis (PLS-DA) in male *16p11.2^df/+^*and WT astrocytes. (D) VIP scores, showing the top 15 (score > 1) metabolite drivers contributing to the separation of metabolic profiles identified by PLS-DA in female *16p11.2^df/+^*and WT astrocytes. (E) *Left,* PLS-DA on metabolomics data from male *16p11.2^df/+^* (n=3) and WT (n=3) astrocytes. PLS-DA was obtained with 2 components. The explained variances are shown in parentheses. *Right,* Volcano plot showing most significantly altered metabolites (p<0.05). Plot summarizes both fold-change and t-test criteria. Scatter-plot of the negative log10-transformed p-values from the t-test plotted against the log2 fold change is shown. (F) *Left,* PLS-DA of metabolomics data from female *16p11.2^df/+^* (n=3) and WT (n=3) astrocytes. PLS-DA was obtained with 2 components. The explained variances are shown in parentheses. *Right,* Volcano plot showing most significantly altered metabolites (p<0.05). Plot summarizes both fold-change and t-test criteria. Scatter-plot of the negative log10-transformed p-values from the t-test plotted against the log2 fold change is shown. (G) Details of most significantly altered metabolites from male *16p11.2^df/+^* (n=3) and WT (n=3) astrocytes. Identified with negative log10-transformed p-values from the t-test plotted against the log2 fold change normalized by Pareto scaling. *Left,* Violin plot of adenosine **(p=0.005) (yellow data point represents mean, black data point represents individual mouse). *Right,* Violin plot of alpha-ketoglutaric acid **(p=0.005) (yellow data point represents mean, black data point represents individual mouse). (H) Details of most significantly altered metabolites from female *16p11.2^df/+^* (n=3) and WT (n=3) astrocytes. Identified with negative log10-transformed p-values from the t-test plotted against the log2 fold change normalized by Pareto scaling. *Left* Violin plot of alpha-ketoglutaric acid ***(p=0.0004) (yellow dot represents mean, black data points represent individual mice). *Right* Violin plot of fructose 1,6-biphosphate *(p=0.012) (yellow dot represents mean, black data points represent individual mice).

A significant decrease in adenosine was noticeable in the male *16p11.2^df/+^* astrocytes compared to WT cells (p<0.005) (Figure 1A,E,G). Adenosine acts as a neuromodulator associated with the initiation of sleep, suggesting a potential link to the sleep disturbances previously reported in male *16p11.2^df/+^* mice^19–21^. Metabolites significantly increased in male *16p11.2^df/+^* astrocytes included alpha-ketoglutaric acid (AKG) (p<0.005), an intermediary of the Tricarboxylic acid (TCA) cycle^22^, N-acetylglutamic acid (p<0.007) involved in brain development, carnosine (p<0.013) which has anti-inflammatory properties^23,24^, orotic acid (p<0.022) required for RNA synthesis^25^, and uridine (p<0.038) which regulates chemokine expression^26^ (Figure S1C). A significant increase in ribose 5-phosphate (p<0.042) was also seen in males *16p11.2^df/+^* astrocytes, suggesting dysregulation of the pentose phosphate pathway (PPP)^27^, as we previously suggested in whole-cortex extracts^13^. As opposed to what was found in males, AKG was significantly reduced (p<0.0004) in *16p11.2^df/+^* female astrocytes compared to WT cells (Figure 1F,H), which could indicate decreased oxidative phosphorylation (OXPHOS)^28,29^. There was also a reduction of N-acetylglutamic acid (p<0.023) and carnosine (p<0.032), opposite to what was seen in male astrocytes (Figure S1D). Additionally, LC-MS detected a reduction in N-acetylaspartic acid (p<0.009) and an increase in Fructose 1,6-biphosphate (p<0.012), an intermediary of glycolysis^30,31^ (Figure 1F,H). Sex specific differences were confirmed by direct comparisons (Figure S2). Surprisingly, alpha-ketoglutaric acid appeared higher in female WT astrocytes (p<0.022), and adenosine lower (p<0.005), compared to WT male counterparts. Additionally, a clear pattern of significantly elevated metabolites was identified in the male *16p11.2^df/+^* astrocytes versus female *16p11.2^df/+^*cells (Figure S2E). Despite the increase in AKG in the WT female astrocytes, male *16p11.2^df/+^*astrocytes still displayed a significant increase in AKG (p<0.0001) (Figure S2H).

Altogether, these results reveal sex-specific changes in metabolite profiles between WT and *16p11.2^df/+^*cultured primary cortical astrocytes.

### Metabolite changes are not reflective of baseline mitochondrial alterations

We next sought to determine if the cellular capacity for energy production was altered in *16p11.2^df/+^* cortical astrocytes. Since intracellular metabolite profiling by LC-MS indicated possible alterations in energy production, we quantified mitochondrial network density by immunocytochemistry, and mitochondrial function using a Seahorse XF Cell Mito Stress Test. Mitochondrial density appeared similar in *16p11.2^df/+^* and WT male derived astrocytes (Figure S3A,B), indicating no change in mitochondrial mass, consistent with our recent findings *in vivo*^13^. The Seahorse test was used to measure Oxygen Consumption Rate (OCR), a proxy to oxidative phosphorylation, and Extracellular Acidification Rate (ECAR), representative of glycolysis. No significant difference was seen for both OCR and ECAR in cortical astrocytes isolated from both male and female *16p11.2^df/+^*and WT littermates at any timepoint in the assay (Figure S3C-F).

These results suggest that 16p11.2 haploinsufficiency does not alter mitochondrial function in primary cortical astrocytes, similar to what we reported in primary cortical endothelial cells from the same mouse model^32^.

### Glucose availability affects mitochondrial function in 16p11.2-deficient astrocytes

Since mitochondrial function appeared unaffected in astrocytes from *16p11.2^df/+^* mice in basal culture conditions, we tested whether metabolic function in these cells was sensitive to energy substrate bioavailability. Our standard Seahorse test was conducted using media with a mid-range glucose concentration (10 mM). However, astrocytes are typically cultured in higher glucose conditions (25 mM). Thus, we challenged the cells with either low (5 mM) or high (25 mM) glucose concentrations in culture for a week prior to the Seahorse assay (Figure 32G,H). An osmotic control (19.5 mM mannitol) was included with the low glucose media. The assay was performed using Seahorse media with corresponding glucose concentrations. In low glucose conditions, we measured reduced OCR at baseline (1.3 min p<0.001, 7.8 min p<0.0001, 14.3 min p<0.0004) in *16p11.2^df/+^* astrocytes compared to cells from WT littermates (Figure S3G). There was no change in glycolysis as suggested by normal ECAR. In high glucose conditions, the OCR was not affected in *16p11.2^df/+^* astrocytes, albeit more variable in WT cells (Figure S3H). However, *16p11.2^df/+^* cortical astrocytes achieved ECAR at later time points (60.3min, p<0.044). These findings suggesting 16p11.2 haploinsufficiency affects astroglial metabolism in a context-dependent manner.

### DNA methylation is differentially regulated in 16p11.2-deficient astrocytes

Mounting evidence shows that regulation of gene expression via epigenetic mechanisms, including DNA methylation, is central in ASD^33–35^. Metabolomics performed with cultured cells extracts revealed a significant increase in alpha-ketoglutaric acid (AKG) in male *16p11.2^df/+^* astrocytes, but a significant decrease in female *16p11.2^df/+^* cells compared to sex-matched WT controls (Figure 1). This may suggest an important role for AKG as a molecular switch in astrocytes from an ASD mouse model. Interestingly, AKG is a known epigenetic modulator acting as cofactor for enzymes involved in DNA demethylation^36^. AKG regulates reprogramming of DNA methylation during early developmental stages^37^, and modulates methylation and hydroxymethylation marks^38^. As ASD are biased towards males and prior work reported mostly male-specific changes in *16p11.2^df/+^*mice^13,20,39,40^, we conducted combined genetic and epigenetic DNA sequencing. Genomic DNA was extracted acutely from primary cortical astrocytes isolated from male *16p11.2^df/+^* and WT littermates and analyzed using a Biomodal^®^ assay to identify differentially hydroxymethylated (DhMRs) and methylated (DMRs) regions. A multiple comparison (false discovery ratio, FDR of 0.1) elicited 287 hydroxymethylation differences and 50,047 methylation differences between *16p11.2^df/+^* and WT astroglial DNA (Figure 2A). Hydroxymethylation activity peaked on chromosome 13, with 77 identified DhMRs (Figure 2B), and methylation activity was concentrated on chromosome 3 with 7,736 DMRs (Figure 2C). While DhMRs and DMRs were evenly dispersed throughout chromosome 3 (Figure 2D), DhMRs on chromosome 13 appeared isolated to a hot spot (Figure 2E). DhMRs on Chromosome 3 were in both intragenic and intergenic, with 16 intragenic and 27 intergenic regions (Figure 3A,B). DhMRs on chromosome 13 appear predominantly intergenic, with only 6 intragenic and 71 intergenic regions. The DMRs on chromosome 3 were divided between intragenic (2,605) and intergenic (4,731). On chromosome 13, there were 754 intragenic DMRs and 1,163 intergenic DMRs (Figure 3A,B).

**Figure 2.**
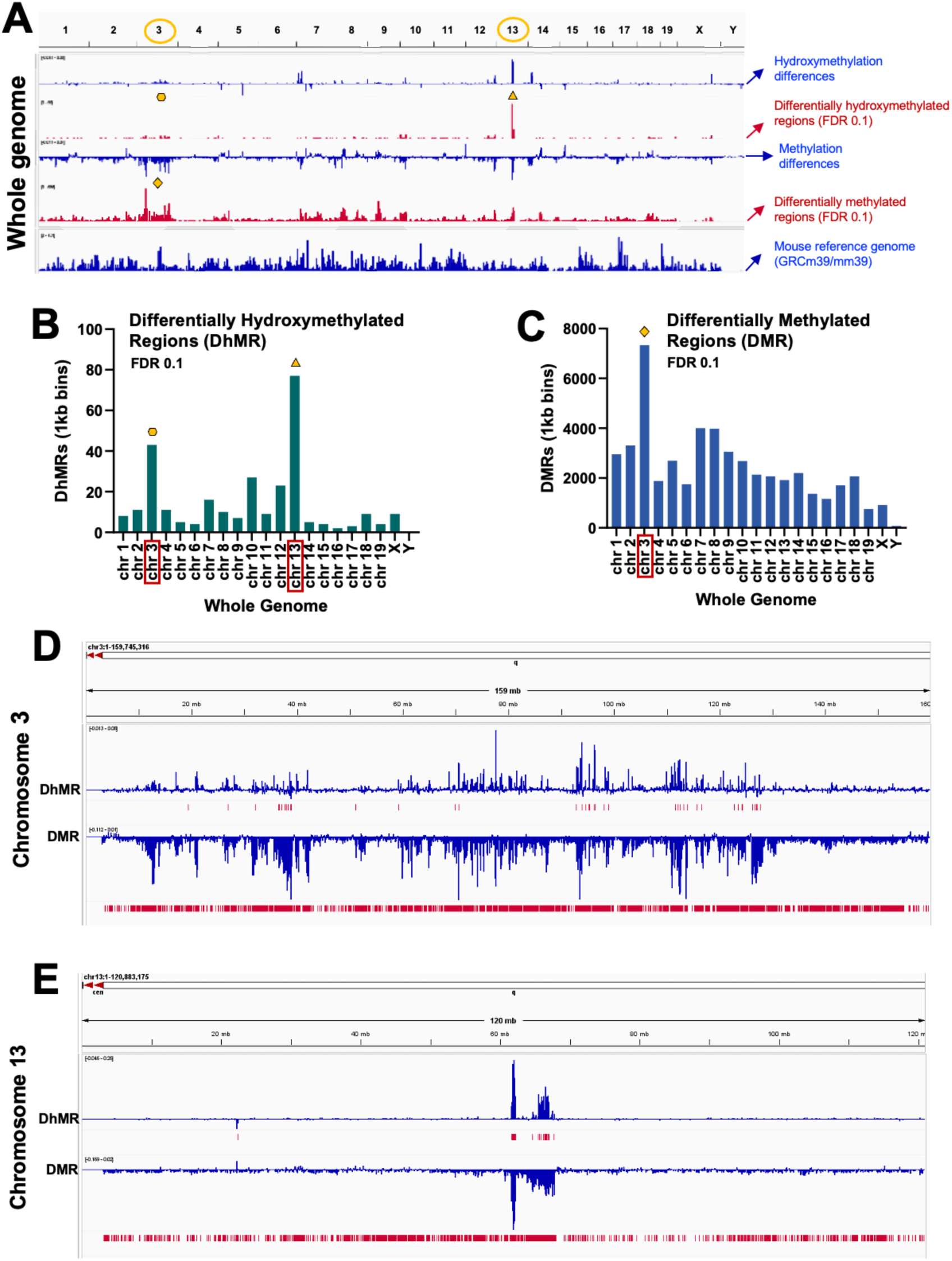
Differential epigenetic modulation assessed in male 16p11.2-deficient P8 astrocytes. (A) Integrative Genomics Viewer (IGV) snapshot of whole-genome sequencing reads from male *16p11.2^df/+^* and WT astroglial genomic DNA purified from 3 mice (n=3) per genotype. Hydroxymethylation and methylation read differences are represented in blue tracks, with Differentially hydroxymethylated Regions (DhMRs) and Differentially Methylated Regions (DMRs) meeting multiple-comparison-corrected p-value (false discovery rate, FDR of 0.1) represented in red marks. Bottom track is mouse reference genome (GRCm39/mm39). Peak genomic activity areas are highlighted with yellow symbols. (B) Chart of identified DhMRs (FDR 0.1) by chromosome. (C) Chart of identified DMRs (FDR 0.1) by chromosome. (D) IGV snapshot detailing DhMR and DMR tracks for chromosome 3 (chr3:1-159,745,316). (E) IGV snapshot detailing DhMR and DMR tracks for chromosome 13 (chr13:1-120,883,175).

**Figure 3.**
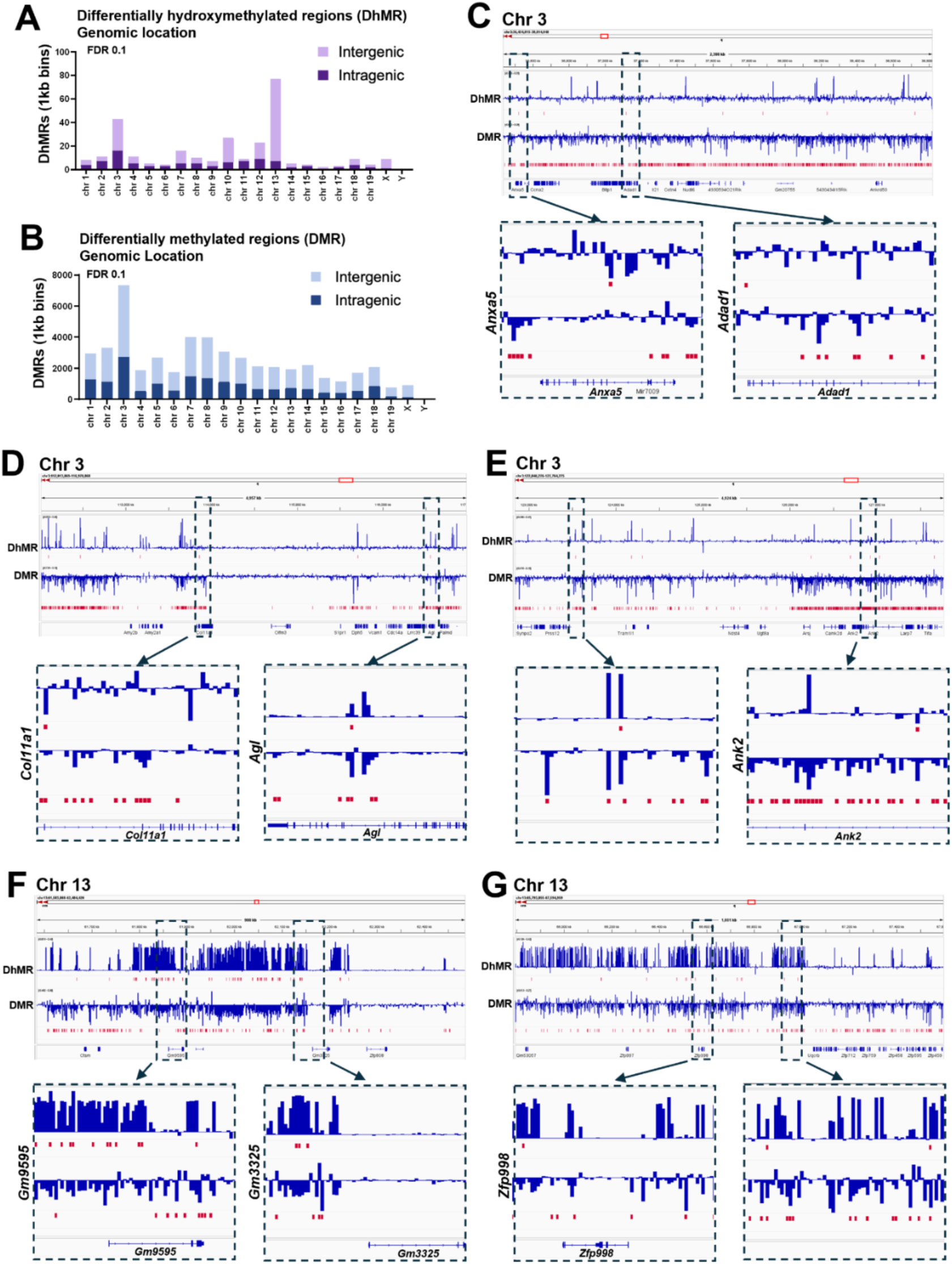
Changes in intragenic and intergenic methylation on chromosomes 3 and 13 from male 16p11.2-deficient P8 astrocytes. (A) Chart outlining whole-genome division in differentially methylated regions (DMRs) between intragenic and intergenic placement. (B) Chart outlining whole genome division in differentially hydroxymethylated regions (DhMRs) between intragenic and intergenic placement. (C) Integrative Genomics Viewer (IGV) snapshot of hydroxymethylation and methylation differences identified on male *16p11.2^df/+^* and WT astroglial genomic DNA purified from 3 mice (n=3) per genotype. Hydroxymethylation and methylation read differences represented in blue tracks. Differentially Hydroxymethylated Regions (DhMRs) and Differentially Methylated Regions (DMRs) meeting multiple-comparison-corrected p-value (false discovery ratio, FDR 0.1) represented in red marks. Bottom track is mouse reference genome (GRCm39/mm39). *Top panel,* IGV snapshot of chromosome 3 region containing increased hydroxymethylation and methylation (chr3:36,434,815-38,814,948). *Bottom panel left,* Detailed view of *Anxa5* intragenic region (chr3:36,445,984-36,594,741). *Bottom panel right,* Detailed view of *Adad1* intragenic region (chr3:37,073,264-37,222,021). (D) *Top panel,* IGV snapshot of chromosome 3 region containing increased hydroxymethylation and methylation (chr3:113,827,658-113,976,415). *Bottom panel left,* Detailed view of *Col11a1* intragenic region (chr3:113,827,658-113,976,415). *Bottom panel right,* Detailed view of *Agl* intragenic region (chr3:116,488,981-116,637,738). (E) *Top panel,* IGV snapshot of chromosome 3 region containing increased hydroxymethylation and methylation activity (chr3:122,840,276-127,764,275). *Bottom panel left,* Detailed view of intergenic region of methylation (chr3:123,462,067-123,610,824). *Bottom panel right,* Detailed view of *Ank2* intragenic region (chr3:126,831,448-126,980,205). (F) *Top panel,* IGV snapshot of chromosome 13 hydroxymethylation hotspot (chr13:61,583,868-62,484,420). *Bottom panel left,* Detailed view of intergenic regions of hydroxymethylation adjacent to *Gm9595* (chr13:61,802,947-61,951,000). *Bottom panel right,* Detailed view of *Gm3325* intragenic region (chr13:62,107,278-62,256,066).(G) *Top panel,* IGV snapshot of chromosome 13 hydroxymethylation hotspot (chr13:65,793,855-67,594,959). *Bottom panel left,* Detailed view of intergenic regions of hydroxymethylation surrounding *Zfp998* (chr13:66,504,209-66,653,000). *Bottom panel right,* Detailed view of intergenic regions of hydroxymethylation (chr13:66,885,144-67,033,651).

On chromosome 3, intragenic regions impacted by hydroxymethylation included the genes *Annexin A5* (*Anxa5*) comprising 1 DhMR and 3 DMRs, and *Adenosine deaminase domain* (*Adad1*) with 1 DhMR and 7 DMRs (Figure 3C). The human ortholog of *Anxa5* is associated with nervous system autoimmune disease^41^. When active, it provides anti-inflammatory protection and was linked to Lewy Body dementia^42^ and amyloid plaque clearing^43^. Regionally found in the brain^44^, *Adad1* is associated with RNA binding^45^. Additionally, *Collagen type XI alpha 1* (*Col11a1*) contains 1 DhMR and 28 DMRs (Figure 3D). Expressed in the CNS^46^, *Col11a1* encodes the alpha 1 chain of type XI collagens and is particularly important during development^47,48^. We also identified 1 DhMR and 10 DMRs within *Amylo-1,6-glucosidase, 4-alpha-glucanotransferase* (*Agl*) (Figure 3D). *Agl* is expressed in the CNS^49,50^ and its deficiency leads to glycogen storage disease^51^. *Agl* encodes for a glycogen debranching enzyme essential for the conversion of glycogen to glucose, without this enzyme glycogenolysis is disrupted^52^. Of note, *Ankyrin 2* (Ank2) contained 2 DhMRs and 198 DMRs (Figure 3E), indicating a high level of methylation and suggesting reduced expression. *Ank2* was previously associated with neural stem cell differentiation and migration^53^. More importantly, *Ank2* has been identified as a high confidence ASD gene^54,55^, suggesting a potential link between 16p11.2 haploinsufficiency and reduced *Ank2* function.

Chromosome 13 presented a narrow area of intense hydroxymethylation that predominantly impacted intergenic regions (Figure 3F,G). Most intragenic DhMRs were found in predicted gene *Gm9595*, additionally containing 7 DMRs (Figure 3F). No gene function has been identified at this time for *Gm9595*, warranting further investigation. *Zinc finger protein 998* (*Zfp998*), which is functionally associated with DNA transcription and RNA metabolic process (GO_REF:0000033; GO_REF:0000033), contained 1 DMR (Figure 3G).

Conversely, the remaining copy of the 16p11.2 (7qF3) locus on chromosome 7, site of the hemideletion, was largely spared DNA methylation changes. Only *Tmem219* and *Mvp* genes showed a restricted DMR (Figure S4).

A Genomics Regions Enrichment of Annotations Tool (GREAT) analysis of identified DhMRs did not reveal significant links to gene ontological functions. However, there were identified interactions related to the observed methylation changes. GO terms pointed to structural changes as well as changes associated with cellular responses and immune functions (Figure S5,D).

Overall, these results demonstrate that male *16p11.2^df/+^* astrocytes display differential DNA methylation, particularly on chromosomes 3 and 13, thus impacting gene expression remotely from the 16p11.2 (7qF3) deletion site.

### Sex-specific gene expression profiles in 16p11.2-deficient cortical astrocytes

As mentioned above, several genes displayed sites of hydroxymethylation or methylation, suggesting that epigenetic modifications in 16p11.2-deficient cortical astrocytes may alter transcriptional activity. Initial metabolomic experiments also established major changes in *16p11.2^df/+^* astrocytes, especially for the epigenetic regulator AKG which appeared opposed in male versus female cells. This led us to assess whether metabolite and epigenetic changes were correlated with transcriptional differences. To this end, we performed bulk sequencing of total RNA acutely extracted from cortical P8 astrocytes from male and female *16p11.2^df/+^*and WT littermates. A total of 20,937 targets were detected from astrocyte-enriched transcripts (Figure S6A).

In male samples, 33 differentially expressed genes (DEGs) were identified (FDR threshold of 0.05), 4 of which were upregulated and 29 downregulated. Of these downregulated genes, 18 related to the 16p11.2 (7qF3) excision locus, as expected, confirming validity of the model (Figure 4A). A cluster of downregulated genes was localized to chromosome 4 (*Arhgef19, Epb41l4b, Fbxo44, Gm29367, Miip, Tnfrsf8, Zfp982, Znf41-ps*) (Figure 4A,B). When considering gene ontological associations for male DEGs, the most common biological process was ‘Response to stimulus’ (GO_REF:0000033, GO_REF:0000119, GO_REF:0000002) evident in 2 upregulated genes (*Ackr3, Ets1*) and in 4 downregulated genes (*Arhgef19, Epb41l4b, Fbxo44, Tnfrsf8*), suggesting changes in the reactive state of male *16p11.2^df/+^* astrocytes (Figure 4C). Several identified DEGs, including 2 upregulated (*Ablim2, Ets1*) and 2 downregulated (*Arhgef19, Epb41l4b*) genes, were associated with ‘cellular component organization’ (GO_REF:0000002, GO_REF:0000008, GO_REF:0000096). Additionally, *Ablim2* upregulated in male *16p11.2^df/+^* astrocytes potentially impacts ‘cytoskeletal protein binding’ (GO_REF:0000002, GO_REF:0000033). Of note, the downregulated genes *Arhgef19* and *Epb41l4b* were associated with cytoskeletal formation and wound healing functions^56^. *Arhgef19* encodes a Guanine nucleotide Exchange Factor (GEF) for Cdc42, inducing changes in cell morphology such as formation of filopodia, stress fibers, and lamellipodia^57^. Interestingly, mutations in another Rho guanine nucleotide exchange factor (RhoGEF) family gene, *ARHGEF9* (X linked in humans) have been linked to ASD and epileptic encephalopathy^58–60^. These transcriptomic changes in male cells potentially signify changes in astroglial morphogenesis, which remains to be thoroughly investigated. In the RNA samples from female astrocytes, a total of 24 DEGs were identified, comprising 3 upregulated and 21 downregulated genes in *16p11.2^df/+^* astrocytes compared to WT cells (Figure S6B). In the case of females, all downregulated genes were associated with the excision locus (Figure S6B). Expression of *Cdk5r2* and *Dnajc6* was upregulated. *Cdk5r2* is associated with nervous system development, and its encoded protein P39 has been tangentially linked with activated astrocytes through the activation of cyclin-dependent kinase 5^61^. *Cdk5r2* is also linked the biological process ‘cytoskeletal protein binding’^62^ (Figure S6C), suggesting possible morphological changes in these cells as well^63^. Both upregulated DEGs are associated with the GO term ‘cellular component organization’^64,65^ as seen in the males (Figures 4C and S5C).

**Figure 4.**
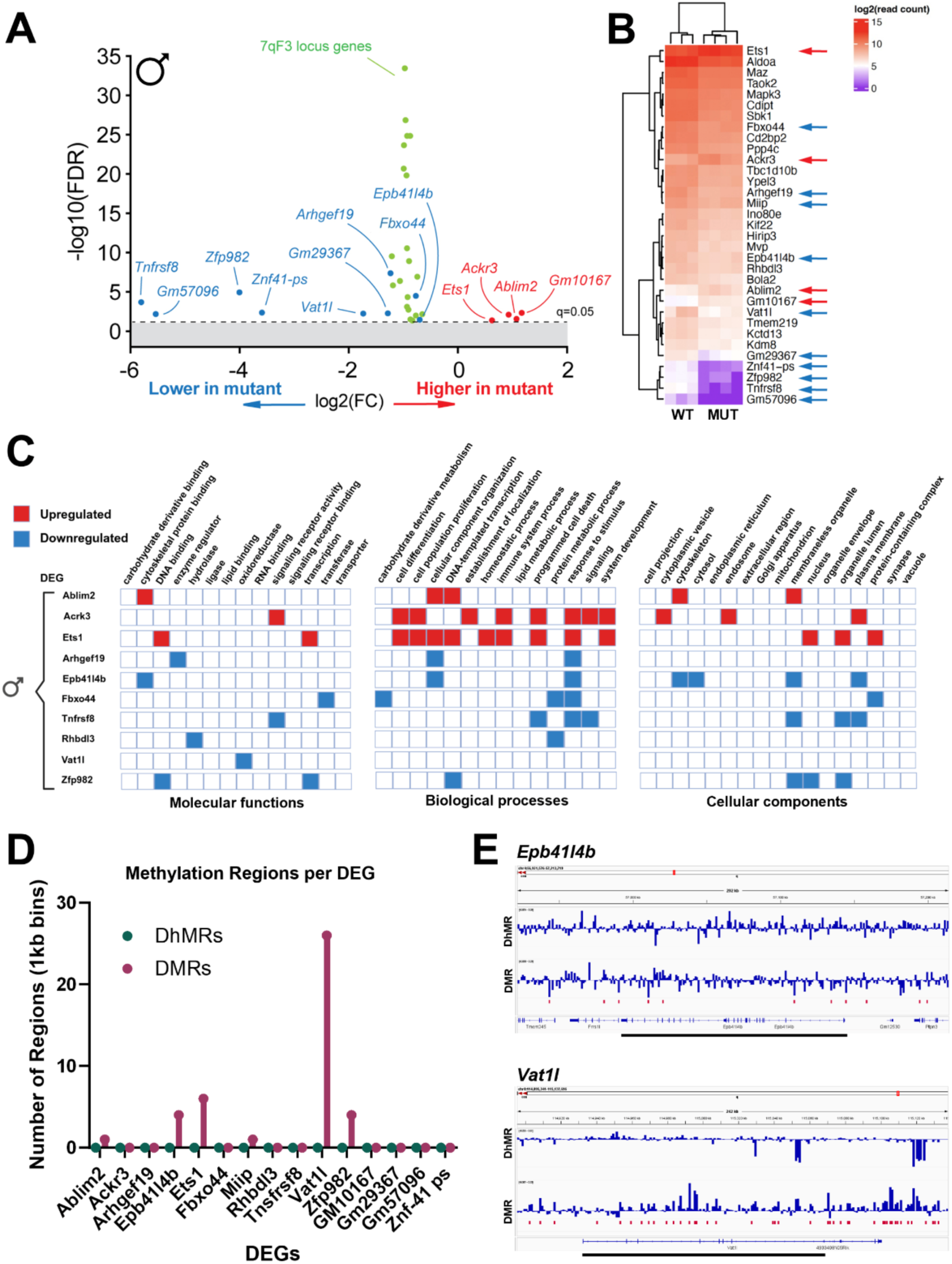
Differential gene expression in male *16p11.2^df/+^* cortical P8 astrocytes. (A) Volcano plot of differentially expressed genes (DEGs) identified with false discovery rate (FDR)-adjusted q-value<0.05 for male *16p11.2^df/+^* (n=4) and WT (n=3) astrocytes (upregulated genes indicated in red, and downregulated genes in blue, 7qF3 locus genes in green). (B) Heatmap (multivariate statistical analysis) representing significant DEGs from male *16p11.2^df/+^* (n=3) and WT (n=4) astrocytes (upregulated genes indicated by red arrows, downregulated genes by blue arrows). (C) Gene Ontology terms associated with identified DEGs (red boxes represent upregulated genes, blue boxes represent downregulated genes). (D) Chart outlining Differentially Hydroxymethylated Regions (DhMRs) and Differentially Methylated Regions (DMRs) classified by DEG. (E) Integrative Genomics Viewer snapshot of hydroxymethylation and methylation differences between male *16p11.2^df/+^* (n=3) and WT (n=3) astroglial genomic DNA related to two downregulated DEGs.

Interestingly, the pseudogene *Gm10167* (MGI:3704263), found on chromosome 5, was upregulated in both male and female *16p11.2^df/+^*astrocytes. Though there is no known function associated with this gene at the moment, its upregulation in both sexes merits future pursuit.

When cross-referencing DEGs identified by RNA sequencing and methylated regions identified by DNA sequencing in male astrocytes, no DhMR was found in any DEG (Figure 4D). However, 26 DMRs were present in *Vat1l*, 6 DMRs in *Ets1*, 4 DMRs in *Zfp982*, and 4 DMRs in *Epb41l4b* (Figures 4E and S7). *Vat1l* found on chromosome 8 and encoding Vesicle Amine Transport Protein 1-like was associated with ‘oxidoreductase activity’ (GO_REF:0000043). *Epb41l4b* found on chromosome 4 and encoding Band 4.1-like protein 4B, was associated with ‘cytoskeletal protein binding’ (GO_REF:0000002). *Ets1* found on chromosome 9 encodes an important skeletogenic transcription factor controlling cellular differentiation, migration and proliferation^66–68^. Finally, *Zfp982* found on chromosome 4 encodes a transcriptional regulator of stemness^69^.

Altogether, these findings reveal distinct, sex-specific transcriptomic profiles in 16p11.2 deficient mouse cortical astrocytes. While commonalities were seen in gene ontology suggesting changes in cytoskeletal remodeling and cellular morphology, these potential changes may not be facilitated through the same pathways in both sexes.

## DISCUSSION

This *in vitro* study establishes metabolic, epigenetic and transcriptomic signatures in primary cortical astrocytes from a 16p11.2 deletion ASD mouse model. We reveal how 16p11.2 haploinsufficiency affects the astroglial metabolome and transcriptome in a sex-dependent manner, and how it leads to DNA methylation changes in male astrocytes, altogether pointing to remote genomic effects of a 16p11.2 (7qF3) locus deletion.

Metabolomics findings support the existence of important sex differences in intracellular metabolite abundance in *16p11.2^df/+^* astrocytes. In particular, AKG was increased in male *16p11.2^df/+^* astrocytes, but decreased in female cells. AKG is an intermediary of the TCA cycle, which could affect oxidative phosphorylation^28^. But AKG is also a cofactor for enzymes involved in DNA demethylation^36^ and regulates reprogramming of DNA methylation during development^37^. AKG was also shown to modulate aberrant gene methylation and hydroxymethylation marks in cardiovascular disease^38^. Thus, AKG may act as a key molecular switch in this context. However, whether AKG directly leads to differential DNA methylation and gene expression in *16p11.2^df/+^* astrocytes remains to be tested. Additionally, the increase in Ribose-5 phosphate in male *16p11.2^df/+^* astrocytes points of a shift towards the Pentose Phosphate Pathway (PPP) and a consequent reduction in active glycolysis. A shift towards the PPP pathway was also suggested by our previous work in whole-cortex extract from *16p11.2^df/+^* male mice^13^. Female-derived astrocytes showed a reduction in AKG and an increase in Fructose 1-6 biphosphate, suggestive of increased glycolysis. However, there was no evident increase in OCR or ECAR in either sex in the Seahorse Mito Stress test in standard glucose conditions. We previously reported that constitutive male *16p11.2^df/+^* mice exhibited elevated cerebral glucose uptake after peripheral glucose injection^13^, suggesting that *16p11.2^df/+^*cortical astrocytes may be functioning in high glucose conditions. Repeating the Seahorse test in varied glucose concentrations elicited different results. The *16p11.2^df/+^* astrocytes in a low glucose condition (with osmotic control) exhibited increased OXPHOS at baseline, but a reduced glycolytic output over time in a high glucose condition. This suggests limited glycolytic capacity of *16p11.2^df/+^* astrocytes in glucose rich conditions.

Though there was no change directly linked to glutamate in the metabolome of male or female *16p11.2^df/+^* astrocytes, we did observe increases in glutamate products in the male mutant cells. Changes in glutamate dynamics are interesting in the context of literature linking reduction in astrocyte glutamate signaling to ASD-associated behaviors^14–16^. We detected increased abundance of N-acetylglutamic acid in male *16p11.2^df/+^* astrocytes. N-acetylglutamic acid is biosynthesized from glutamate, and its elevation is indicative of increased glutamate uptake. Female *16p11.2^df/+^*astrocytes conversely showed reduced N-acetylglutamic acid. Additionally, we found an increase in carnosine in male *16p11.2^df/+^* astrocytes and reduced in female cells. Carnosine upregulates glutamate transporter GLT-1 in astrocytes, increasing their capacity for clearing glutamate^23^ and reducing inflammation^24^, with anti-epileptic properties^70^. Furthermore, elevated glutamine was found in *16p11.2^df/+^* male astrocytes, suggesting higher capacity to clear glutamate. Excess astrocytic glutamate can also be converted to AKG. The increased abundance of AKG may also account for differential methylation evident in *16p11.2^df/+^* male astrocytes^22^. Of note, the drastic reduction of adenosine in *16p11.2^df/+^* male astrocytes may potentially be associated with sleep-wake cycle alterations found in *16p11.2^df/+^* male mice^20^, that exhibited greater time awake and reduced time in non-rapid eye movement (NREM) sleep. Indeed, adenosine accumulation within the brain’s extracellular fluid during wakefulness is key to promote sleep initiation^71^. Using a mouse model with an astrocyte selective dnSNARE conditionally expressed transgene, blocking astrocyte transmitter release, reduced both total sleep time and NREM duration in a recovery period following sleep deprivation^21^. This phenotype was conserved in WT mice treated intracererbroventricular (i.c.v.) with adenosine A1 receptor antagonist 8-cyclopentyl-1,3-dimethylxanthine (CPT)^21^.

With over 50,047 DMRs in the whole genome, the elevated levels of methylation in male *16p11.2^df/+^* astrocytes underscore significant epigenetic modulation in this glial cell type. *Ank2,* which exhibited a significant amount of methylation with 198 DMRs, is a known ASD risk gene. Yet, an interaction with the 16p11.2 deletion has not been previously identified. *Ank2* is associated with synaptic development, and its methylation in astrocytes may impact synaptic formation, as synaptogenesis mediated by astrocytes is active in this timeframe (P8), reaching it’s peak at P14. Since there is phenotypic overlap between *16p11.2^df/+^ and Ank2* associated ASD, particularly for repetitive behaviours, social deficits and increased rates of seizures, this merits further investigation.

Bulk RNA sequencing further illustrated sex differences, with greater downregulation of genes in *16p11.2^df/+^* male astrocytes. Genes that were upregulated in males or females, though disparate, were associated with similar biological processes, particularly cytoskeletal regulation. *Cdk5r2* encodes protein CDK5 subunit p39 and is known to increase postnatally^72^, recruit scaffolding protein muskelin to stress fibers^73^, and to regulate stress fiber formation and actin organization. *Ablim2* was reported to increase in retinal astrocytes during development^74^, and to stabilize actin by interacting with (pro)renin receptor^75^, potentially reducing cell motility. Furthermore, a GREAT analysis performed on our methylation dataset supports increased methylation in genes associated with developmental structural changes.

Data presented in this study are in line with well-established sex differences in ASD phenotypical expression^20,60,76–78^. Moreover, gene expression studies in mice have found that male astrocytes reach maturity at an earlier timepoint than female astrocytes^79^. Early changes in gene expression in astrocytes at key periods of maturation may account for regional subpopulations and morphological heterogeneity found between males and females^80–82^.

Overall, this study creates a framework upon which future work can expand upon to examine the downstream consequences of these molecular variations beyond development.

## ACKNOWLEDGEMENTS & FUNDING

We thank Drs. Julie Ouellette and Moises Freitas-Andrade (Lacoste lab) for technical assistance. We thank Dr. Mary-Ellen Harper at the University of Ottawa for sharing their SeaHorse analyzer. Metabolites were analysed at the University of Ottawa Metabolomics Core Facility; this facility is supported by the Terry Fox Foundation and Ottawa University. We would like to acknowledge the assistance of StemCore Laboratories Genomics Core Facility (OHRI, uOttawa), RRID:SCR_012601. B.L. is supported by start-up funds from the Ottawa Hospital Research Institute, two *Canadian Institutes of Health Research* project grants (388805, 506513), and an Autism Research Program Idea Development Award from the US Department of Defense (US DoD) office of the Congressionally Directed Medical Research Programs (W81XWH2210583). N.B. was also supported by a doctoral training award from the Brain-Heart Interconnectome at University of Ottawa.

## STAR METHODS

### RESOURCE AVAILABILITY

#### Lead Contact

Dr. Baptiste Lacoste

Faculty of Medicine, Department of Cellular and Molecular Medicine, University of Ottawa.

451 Smyth Road, Ottawa (ON) K1H 8M5, Canada

#### Materials Availability

This study did not generate new unique reagents.

#### Data and Code Availability

All data reported in this paper will be shared by the lead contact upon request.

This paper does not report original code.

Any additional information required to reanalyze the data reported in this paper is available from the lead contact upon request.

### EXPERIMENTAL MODEL AND SUBJECT DETAILS

B6129S-Del(7Slx1b-Sept1)4Aam/J (https://www.jax.org/strain/013128) procured from JAX (Jackson laboratory, stock# 013128). Housed and treated according to protocols approved by the University of Ottawa Animal Care Committee and meeting standards in accordance with the Canadian Council on Animal Care standards. Male *16p11.2^df+/-^* were bred with B6 wild type (WT) females (Jackson laboratory, stock# 101043), producing *16p11.2^df/+^* heterozygotes and WT mice. As advised by Jackson labs, breeding cages were supplied with supplemented breeding chow (2019, Envigo Teklad). Additionally, upon birth of a litter, cages are supplied with Diet gel (76A, ClearH2O). Experiments were performed on male and female, *16p11.2^df/+^* mice on postnatal day 8 (P8). Both mutation and sex were initially assessed visually then confirmed with PCR.

Genotyping as performed for *16p11.2^df/+^* and WT mice using the following primers with a PCR product of 2.2kb for *16p11.2^df/+^* mice^83^:

Forward 5’-CCTCATGGACTAATTATGGAC-3’

Reverse 5’-CCAGTTTCACTAATGACACA-3’

Sex determined by genotyping for Rbm31x/y genes using the following Primers^84^:

Forward 5’-CACCTTAAGAACAAGCCAATACA-3’

Reverse 5’-GGCTTGTCCTGAAAACATTTGG-3’

X chromosome produces 269-bp product and Y chromosome produces 353-bp product.

### METHOD DETAILS

#### Primary mouse brain astrocyte isolation

*16p11.2^df/+^ /B6* mice are euthanized on postnatal day (P)8 by decapitation without anesthesia. Cortices are isolated using autoclaved tools submerged in 100% ethanol for 30 minutes prior dissections. Cortices are placed in DMEM high glucose (Cytiva – High Clone, cat# SH40007.01) and kept over ice. Cortices are manually dissociated by passage through increasing gauge of needle (18g, 27g, and 30g). Homogenized cell slurry is then passed through a 70 μM filter and at 300 g for 10 minutes. Supernatant is aspirated and cell pellet is resuspended in 2 ml of Astrocyte Media [(DMEM HG (Cytiva – High Clone, cat# SH40007.01), 10% Fetal Bovine Serum (FBS) (Wisent, cat# 080-150), and 1% Penicillin/streptomycin (Thermo Fisher Scientific, cat# 15140122)]. Cell suspension is plated in a 6-well plate coated with 0.1% poly-l-lysine (Sigma-Aldrich, cat# 25988-63-0) diluted to 0.01% in sterile MilliQ H_2_O with 2 wells per animal in 2 ml of astrocyte media. Cells are kept in culture at 37 degrees with 5% CO2, with media change every 3 days up to required confluency between 7-10 days.

#### Immunocytochemistry

Primary cortical murine astrocytes were isolated and cultured in Astrocyte Media to 80% confluence (5-7 days post isolation). Upon reaching 80% confluency, cells were passaged and seeded with reduced volume on glass coverslips coated in 0.1% poly-l-lysine (Sigma-Aldrich, cat# 25988-63-0) diluted to 0.01% in sterile MilliQ H_2_O in a 12-well culture plate. Each well received 1 ml of astrocyte media. When desired confluency was reached, cells were fixed with 4% paraformaldehyde (PFA) for 10 mins. Fixed cells were permeabilized in a blocking solution containing 10% donkey serum, 0.1% PBT and 0.5% cold water fish skin gelatin. Astrocytes were stained with guinea pig anti-GFAP (Synaptic Systems, 173004, 1:1000) as a cytoskeletal astrocyte marker, and rabbit anti-SOX9 (Abcam, 1:1000, ab185966) as an astrocyte nuclear marker. Primary antibodies were diluted in blocking solution and incubated for 2 hours at room temperature. Primary antibodies were aspirated and wells are washed with 0.2% PBT. Secondary antibodies (species specific AlexaFluor) were diluted in blocking solution and incubated for 1 hour at room temperature. Wells were washed with 0.2% PBT and then 0.1 M PB. Glass cover slips were adhered to slides using Fluromount G and imaged with Zeiss Axio Imager M2 microscope equipped with digital camera (Axiocam 506 mono) and Zeiss ApoTome.2 module using x10, and x20 objectives. Images were processed with FIJI ImageJ.

#### Metabolomics

Extracted astrocytes were cultured to confluency of 80% then split from 2 to 7 wells in 6-well cell culture plates, 6 wells were used for experimental assay and the remaining well was used for cell counting for result normalization. Astrocytes were then cultured to 80-90% confluency and counted. A minimum of 1.5-2 million cells is required for this assay. Cells received 3 washes of 1 ml ammonium formate to arrest cellular metabolism. Cells were collected from plates in 80uL of ice cold methanol:water (1:1) and stored at -80 in 2 ml snap-lock tubes with 6 ceramic beads (1.4 mm, OMNI international, cat# 19-645) per sample. To lyse the cells, 220 µL of acetonitrile was added and repeated bead beating was conducted at 2000 rpm for 60s. Cell mixture that resulted was incubated with dichloromethane:water 2:1 solution and then separated into layers through centrifugation at 4000g for 10 mins. Water soluble metabolites separated into the upper polar phase (aqueous phase), cell debris settled in interphase, and non-polar metabolites were contained in the lower dichloromethane phase (organic phase). From the upper polar phase, 600 μl was extracted for metabolite sequencing using liquid chromatography-mass spectrometry (LC-MS). The upper polar phase was dried using a refrigerated CentriVap Vacuum Concentrator at −4°C (LabConco Corporation, Kansas City, MO). Samples were resuspended in water and run on an Agilent 6470A tandem quadruple mass spectrometer equipped with a 1290 Infinity II ultra-high performance LC (Agilent Technologies) utilizing the Metabolomics Dynamic MRM Database and Method (Agilent), which uses an ion-pairing reverse phase chromatography (https://www.agilent.com/cs/library/applications/5991-8073EN.pdf). This method was further optimized for phosphate-containing metabolites with the addition of 5 µM InfinityLab deactivator (Agilent) to mobile phases A and B, which requires decreasing the backflush acetonitrile to 90%. Multiple reaction monitoring (MRM) transitions were optimized using authentic standards and quality control samples. Metabolites were quantified by integrating the area under the curve of each compound using external standard calibration curves with Mass Hunter Quant (Agilent). No corrections for ion suppression or enhancement were performed, as such, uncorrected metabolite concentrations are presented. Statistical analysis was performed by one-way analysis of variance (ANOVA) followed by Tukey’s post hoc test. P values <.05 were considered statistically significant. Data for group comparisons are presented as box and whisker plots and statistical analysis was performed by an unpaired t test. The box represents the 25-75 interquartile range, and the horizontal line represents the median value. The box plot analysis was performed using Prism 7 (GraphPad Software, La Jolla, CA, USA). Multivariate statistical modelling was performed on log-transformed, mean-centered and pareto-scaled data using MetaboAnalyst 6.0 (https://www.metaboanalyst.ca). Before multivariate analysis, missing values were replaced with 1/3 of the limit of detection of each metabolite. Principal component analysis (PCA) and partial least squares discriminant analysis (PLS-DA) were performed on the identified metabolites. The variables highly contributing to the group separation were selected with a variable importance in projection (VIP)≥1. The clustering analysis and pathway analysis were performed as well using MetaboAnalyst 60.

#### Mitochondrial Density

Astrocytes were isolated from P8 male *16p11.2^df/+^/B6* mice as previously described. Fixed cells were permeabilized in a blocking solution containing 10% donkey serum, 0.1% PBT and 0.5% cold water fish skin gelatin. Astrocytes were stained with guinea pig anti-GFAP (Synaptic Systems, 173004, 1:1000) as a cytoskeletal astrocyte marker, and rabbit anti-TOM20 (Proteintech, 11802-1-AP, 1:200) as a mitochondrial marker. Primary antibodies were diluted in blocking solution and incubated for 2 hours at room temperature. Primary antibodies were aspirated and wells are washed with 0.2% PBT. Secondary antibodies (species specific AlexaFluor) were diluted in blocking solution and incubated for 1 hour at room temperature. Wells were washed with 0.2% PBT and then 0.1 M PB. Glass cover slips were adhered to slides using Fluromount G and imaged with Zeiss Axio Imager M2 microscope equipped with digital camera (Axiocam 506 mono) and Zeiss ApoTome.2 module using x10, and x20 objectives. Images were processed with FIJI ImageJ.Images were forwarded to collaborators at Federal University of São Carlos. The adaptive thresholding algorithm from the Pyvane^85^ library was applied to the TOM20 channel of each sample. First, a Gaussian filter with a standard deviation σ=65 was employed. This parameter, which determines the neighborhood radius for classification, was selected via visual inspection. The filtered image was then subtracted from the original TOM20 channel. Pixels retaining a value exceeding 2.5 were classified as foreground (mitochondria). This threshold was established based on visual inspection. Target cells were manually segmented from the GFAP channel using the GIMP (https://www.gimp.org/) image editor to generate cell masks. Mitochondrial density was calculated as the ratio of the segmented mitochondrial area to the total area of the cell masks.

#### Seahorse XF Cell Mito Stress Test

Astrocytes were cultured to 80-90% confluency then passaged and plated in a Seahorse XF94 microplate (Agilent Technologies, cat# 103794-100) at a density of 20,000 cells per well coated with 0.1% poly-l-lysine (Sigma-Aldrich, cat# 25988-63-0) diluted to 0.01% in sterile MilliQ H_2_O, 80 μl of astrocyte media [(DMEM HG (Cytiva – High Clone, cat# SH40007.01), 10% Fetal Bovine Serum (FBS) (Wisent, cat# 080-150), and 1% Penicillin/streptomycin (Thermo Fisher Scientific, cat# 15140122)] was added to each well. Each sample was plated with a minimum of 3 replicate wells. Sensor cartridge was submerged in XF calibrant in utility plate with attached hydro-booster and placed in non-CO2 37C incubator overnight. After 24 hours media was replaced with Seahorse assay media (Agilent Technologies, cat# 103575-100) with added glucose 10mM (dextrose) (Sigma, cat# D9434-250G), 1mM pyruvate (Sigma-Aldrich, cat# P2256-25G) 2mM l-glutamine (Sigma, cat# G3126-100G) and placed for 1 hour in a 37C-degree incubator with no CO2. Assay was then performed as directed by Agilent Seahorse XF96 Cell Mito Stress Test protocol conducted with Seahorse XF96 Analyzer. Sensor cartridge plate was prepared with assay compounds Oligomycin (1.5 μM), FCCP (1.0 μM), and Rotenone/ Antimycin A (0.5 μM). Sensor cartridge was placed in Seahorse XF96 Analyzer for calibration. Cell plate was then added and baseline function reading was recorded. Cells were selectively injected with loaded compounds for functional assessment of oxygen consumption rate (OCR) and extracellular acidification rate (ECAR): Oligomycin (1.5 μM) at 20 mins, FCCP (1.0 μM) at 40 mins, and Rotenone/ Antimycin A (0.5 μM) at 60 mins. Following the assay, cells were fixed in 4% PFA and stained with DAPI for cell counting using EVOS microscope, verified cell count was used for normalization parameters. Measurement results were reported to Agilent Seahorse Wave Software and exported to Prism 10 (GraphPad Software, La Jolla, CA, USA) for analysis with 2-way ANOVA using Sidak multiple comparison. P values <0.05 were considered statistically significant.

#### Seahorse XF Cell Mito Stress Test with Glucose Challenge

Astrocytes were cultured to 70% confluency then passaged from 2 wells to 3 wells in a 6-well plate. Astrocytes were then cultured in media with either high glucose (25 mM) or low glucose with osmotic control (5.5 mM) with mannitol (19.5 mM) to confluency. Each sample was plated with a minimum of 3 replicate wells. Sensor cartridge was submerged in XF calibrant in utility plate with attached hydro-booster and placed in non-CO2 37C incubator overnight. After 24 hours media was replaced with Seahorse assay media (Agilent Technologies, cat# 103575-100) with added glucose 25mM (dextrose) (Sigma, cat# D9434-250G) or 5.5mM (dextrose) (Sigma, cat# D9434-250G) with 19.5mM of D-mannitol (Sigma-Aldrich, cat# M4125), 1mM pyruvate (Sigma-Aldrich, cat# P2256-25G) 2mM l-glutamine (Sigma, cat# G3126-100G) and placed for 1 hour in a 37C-degree incubator with no CO2. Assay was then performed as directed by Agilent Seahorse XF96 Cell Mito Stress Test protocol conducted with Seahorse XF96 Analyzer. Sensor cartridge plate was prepared with assay compounds Oligomycin (1.5 μM), FCCP (1.0 μM), and Rotenone/ Antimycin A (0.5 μM). Sensor cartridge was placed in Seahorse XF96 Analyzer for calibration. Cell plate was then added and baseline function reading was recorded. Cells were selectively injected with loaded compounds for functional assessment of oxygen consumption rate (OCR) and extracellular acidification rate (ECAR): Oligomycin (1.5 μM) at 20 mins, FCCP (1.0 μM) at 40 mins, and Rotenone/ Antimycin A (0.5 μM) at 60 mins. Following the assay, cells were fixed in 4% PFA and stained with DAPI for cell counting using EVOS microscope, verified cell count was used for normalization parameters. Measurement results were reported to Agilent Seahorse Wave Software and exported to Prism 10 (GraphPad Software, La Jolla, CA, USA) for analysis with 2-way ANOVA using Sidak multiple comparison. P values <0.05 were considered statistically significant.

#### DNA extraction and epigenetic sequencing

*16p11.2^df/+^* /*B6* mice were euthanized at P8 by decapitation without anesthesia. Cortices were separated and hippocampus were removed. Cortices were cut into small pieces then placed in cold HBSS without calcium and magnesium over ice. Using the Miltenyi Neural Dissociation kit (Miltenyi Biotec, cat# 130-092-628) cortices were dissociated then astrocytes were isolated using the MACS magnetic separation with two negative selections and one positive selection. Myelin was isolated and discarded with Myelin removal microbeads (Miltenyi Biotec, cat# 130-096-733). Macrophages were removed with CD45 microbeads (Miltenyi Biotec, cat# 130-052-301). Astrocytes were positively selected with ACSA-2 microbeads (Miltenyi Biotec, cat# 130-097-679). DNA was extracted using Mag-Bind Blood & Tissue DNA HDQ 96 kit (Omega-Biotek, M6399-00). Epigenetic sequencing performed on a Novaseq X (Illumina) platform using 2 × 150 bp paired end reads by OHRI StemCore Laboratories. Libraries were prepared using the Duet EvoC #6101 v4 Library Preparation Kit (Biomodal) with 150 ng of input gDNA according to the manufacturer’s instructions. Hydroxymethylation changes and methylation changes were identified in 1 kilobase bins. Data analysis performed by Ottawa Bioinformatics Core. Visualization images created with Integrative Genomics Viewer^85^.

#### RNA extraction and sequencing

*16p11.2^df/+^* /*B6* mice are euthanized at P8 by decapitation without anesthesia. Cortices were separated and hippocampus were removed. Cortices were cut into small pieces then placed in DMEM high glucose (Cytiva – High Clone, cat# SH40007.01) over ice. Using the Miltenyi Neural Dissociation kit (Miltenyi Biotec, cat# 130-092-628) cortices were dissociated then astrocytes were isolated using the MACS magnetic separation with two negative selections and one positive selection. Myelin was isolated and discarded with Myelin removal microbeads (Miltenyi Biotec, cat# 130-096-733). Macrophages were removed with CD45 microbeads (Miltenyi Biotec, cat# 130-052-301). Astrocytes were positively selected with ACSA-2 microbeads (Miltenyi Biotec, cat# 130-097-679). Isolated astrocytes were pelleted and resuspended with 300 μl of Trizol (ambion, cat# 15596026) to extract RNA. Samples were then stored at -80-degrees until all samples were ready. On the day of RNA extraction, samples were thawed over ice and 300 μl of 100% ethanol was mixed with the trizol astrocyte suspension. RNA from isolated astrocytes was purified with the Direct-zol RNA MicroPrep kit (ZYMO RESEARCH, cat# R2060). Samples were transferred to NRC collaborator for RNA sequencing. RNA sequencing was performed with NextSeq 500/550 High Output Kit v2.5 (150 Cycles) (cat# 20024907) and RNA-seq library was generated with NEBNext Ultra II Directional RNA Library Prep Kit for Illumina. The RNA-seq data were processed by trimming the adaptor sequences, filtering low-quality reads (Phred Score <= 20) and eliminating short reads (length <= 20bps) using software package FASTX toolkit [http://hannonlab.cshl.edu/fastx_toolkit/]). Then the Mouse reference genome (GRCm39 v35)^86^ and corresponding annotation were used as reference for RNA-seq alignment process using STAR (v2.7.11a)^87^. DESeq2^88^ was used for data normalization and differentially expressed gene identification.

A False Discovery Rate (FDR)-adjusted q-value<0.05 was used to identify differentially expressed genes (DEGs).

## DATA AVAILABILITY

Bulk RNA-seq data are accessible through GEO via accession number (GSExxxxxx).

## KEY RESOURCE TABLE

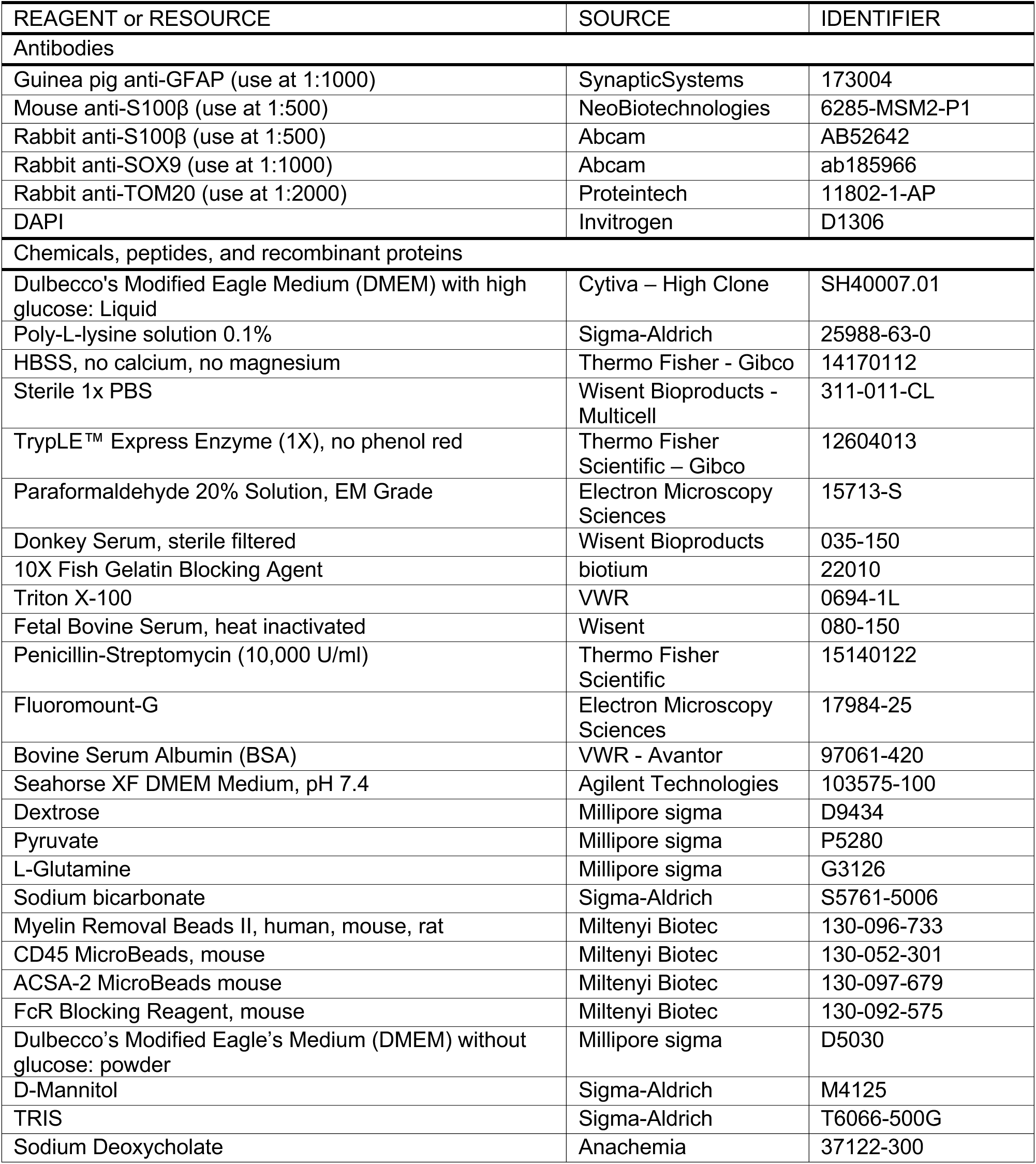
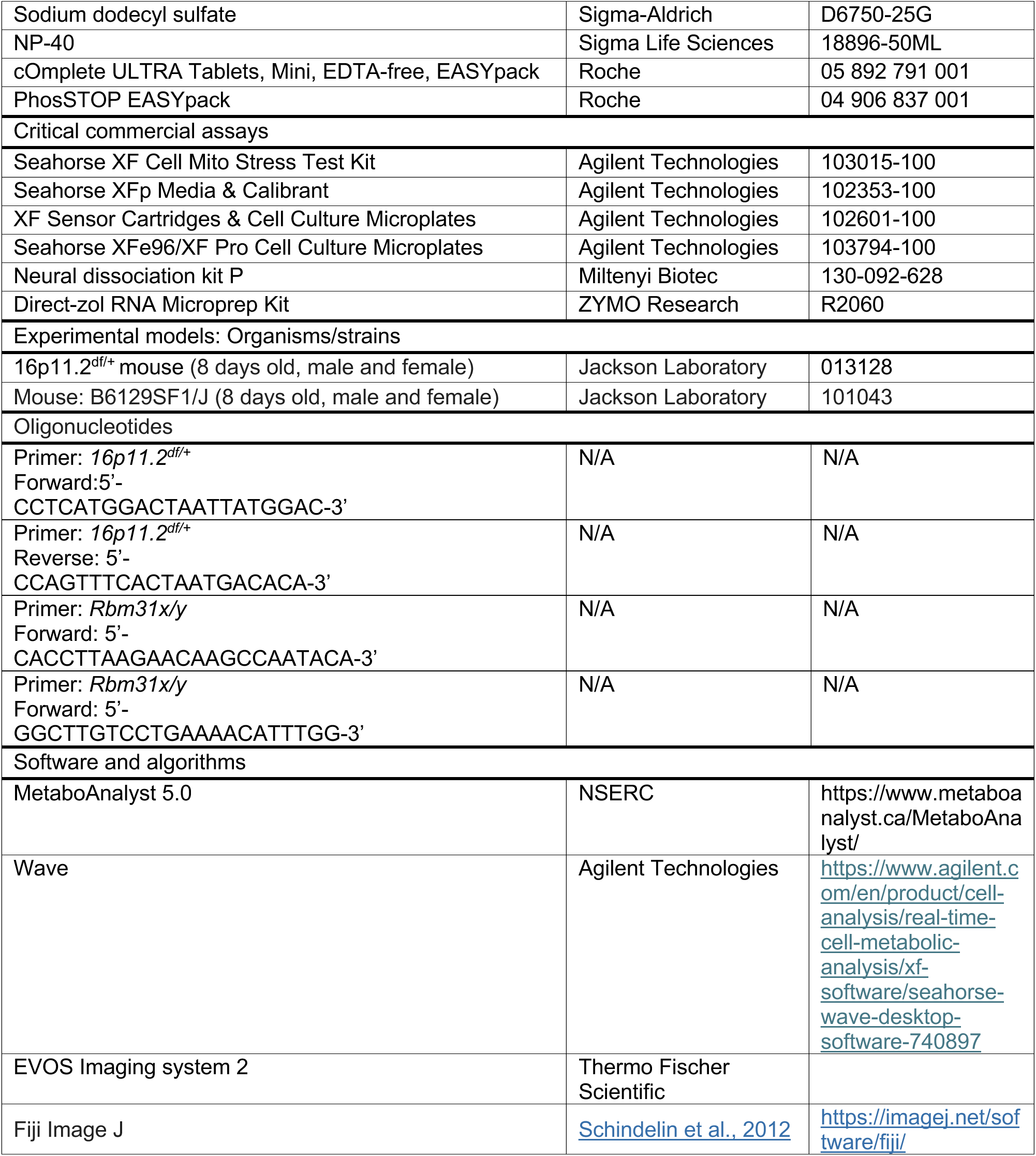
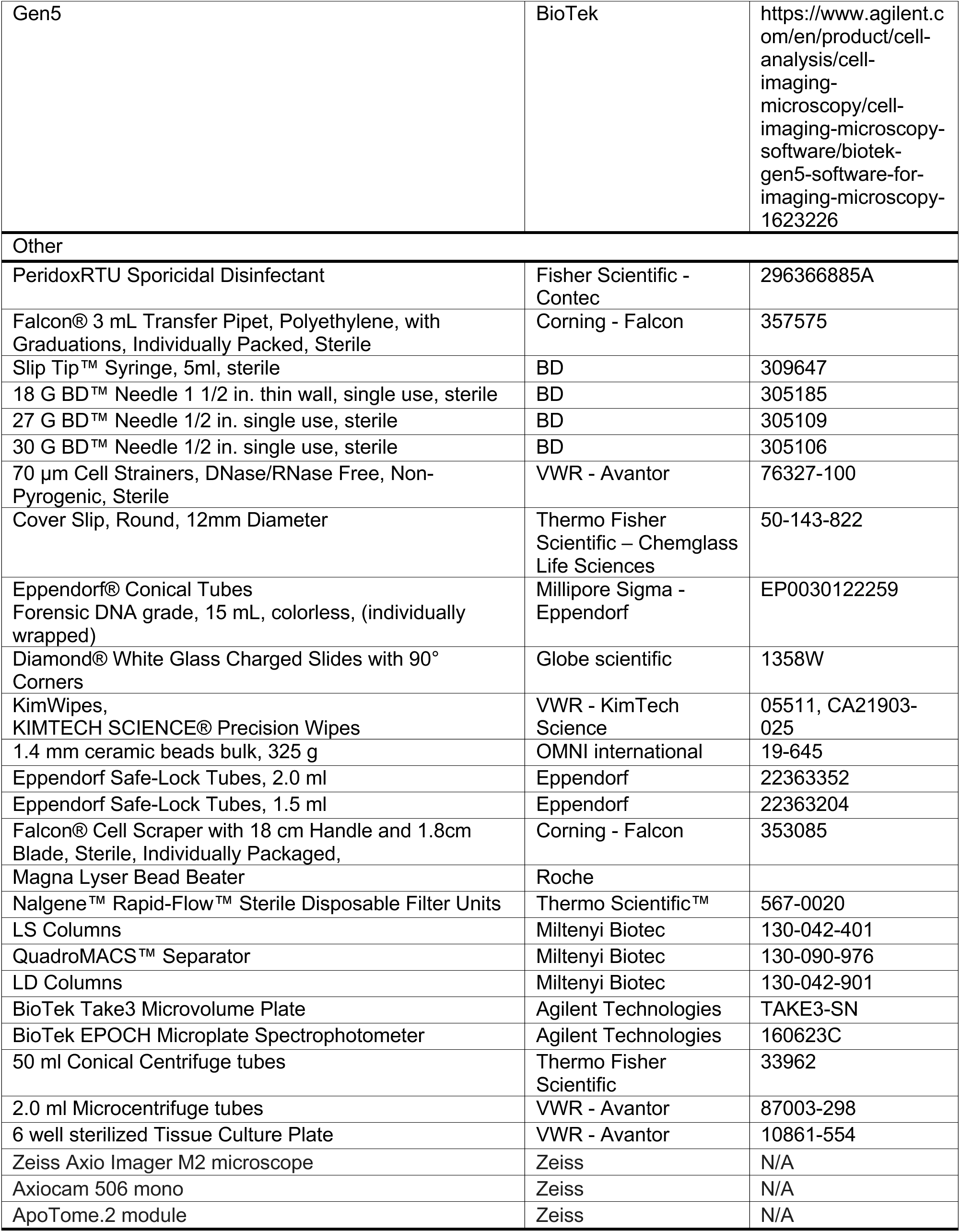

## Supplemental Information

**Figure S1.**
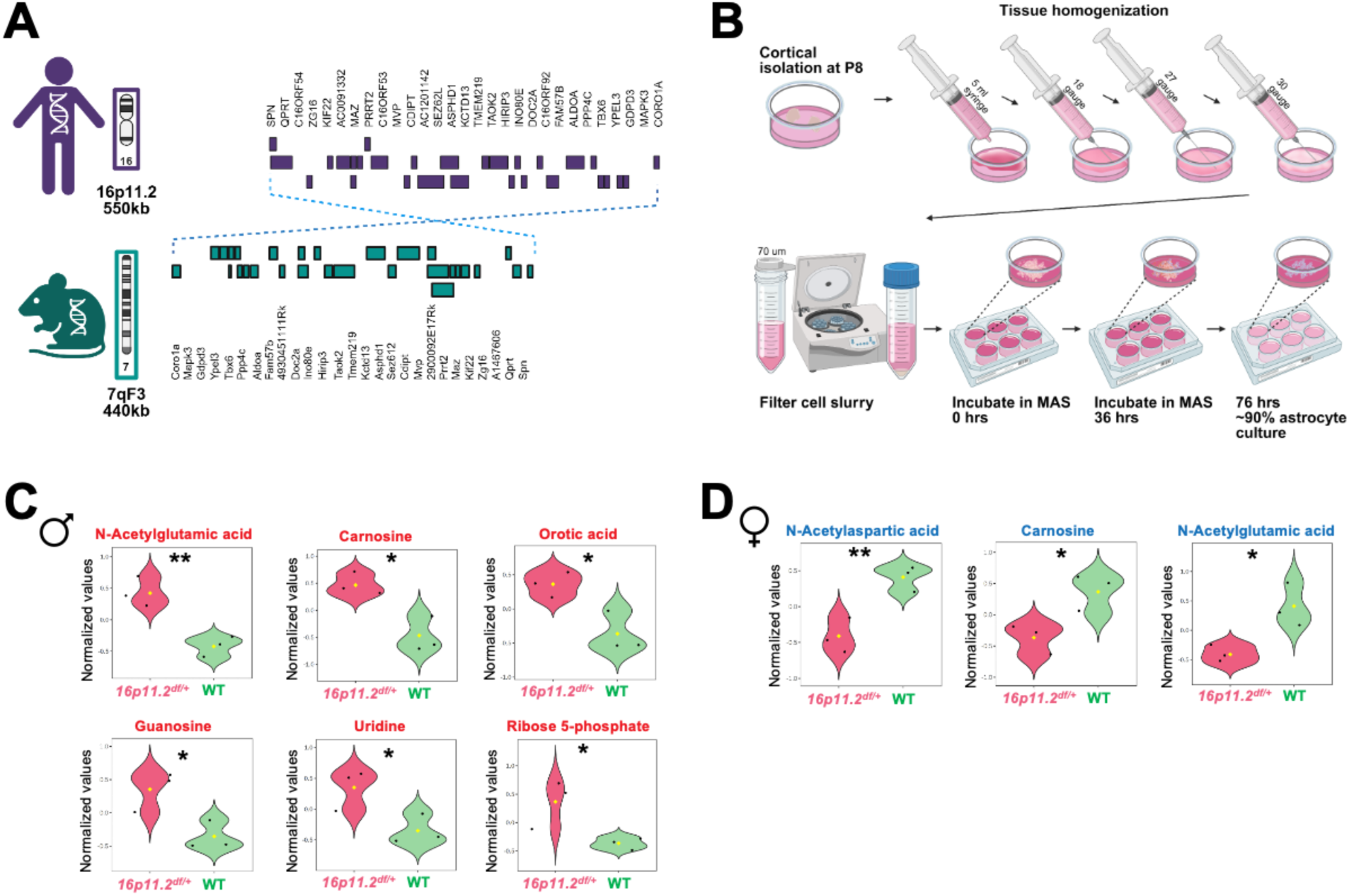
Models and additional information on metabolomics. (A) Schema detailing the human genetic 16p11.2 locus versus murine region of syntenic conservation (7qF3 locus). (B) Graphical summary of cortical astrocyte isolation from mouse pups at postnatal day (P)8. (C) Additional details on most significantly altered metabolites from male *16p11.2^df/+^* (n=3) and WT (n=3) astrocytes, identified with negative log10-transformed p-values from the t-test plotted against log2 fold change normalized by Pareto scaling. Violin plots are shown for N-acetylglutamic acid **(p=0.007), carnosine *(p=0.013), orotic acid *(p=0.022), guanosine *(p=0.030), and uridine *(p=0.038335) (yellow dot represents mean, black data points represent individual mice/samples).(D) Additional details on most significantly altered metabolites from female *16p11.2^df/+^* (n=3) and WT (n=3) astrocytes, identified with negative log10-transformed p-values from the t-test plotted against log2 fold change normalized by Pareto scaling. Violin plots of N-acetylaspartic acid **(p=0.009), carnosine *(p=0.032) and N-acetylglutamic acid *(p=0.023).

**Figure S2.**
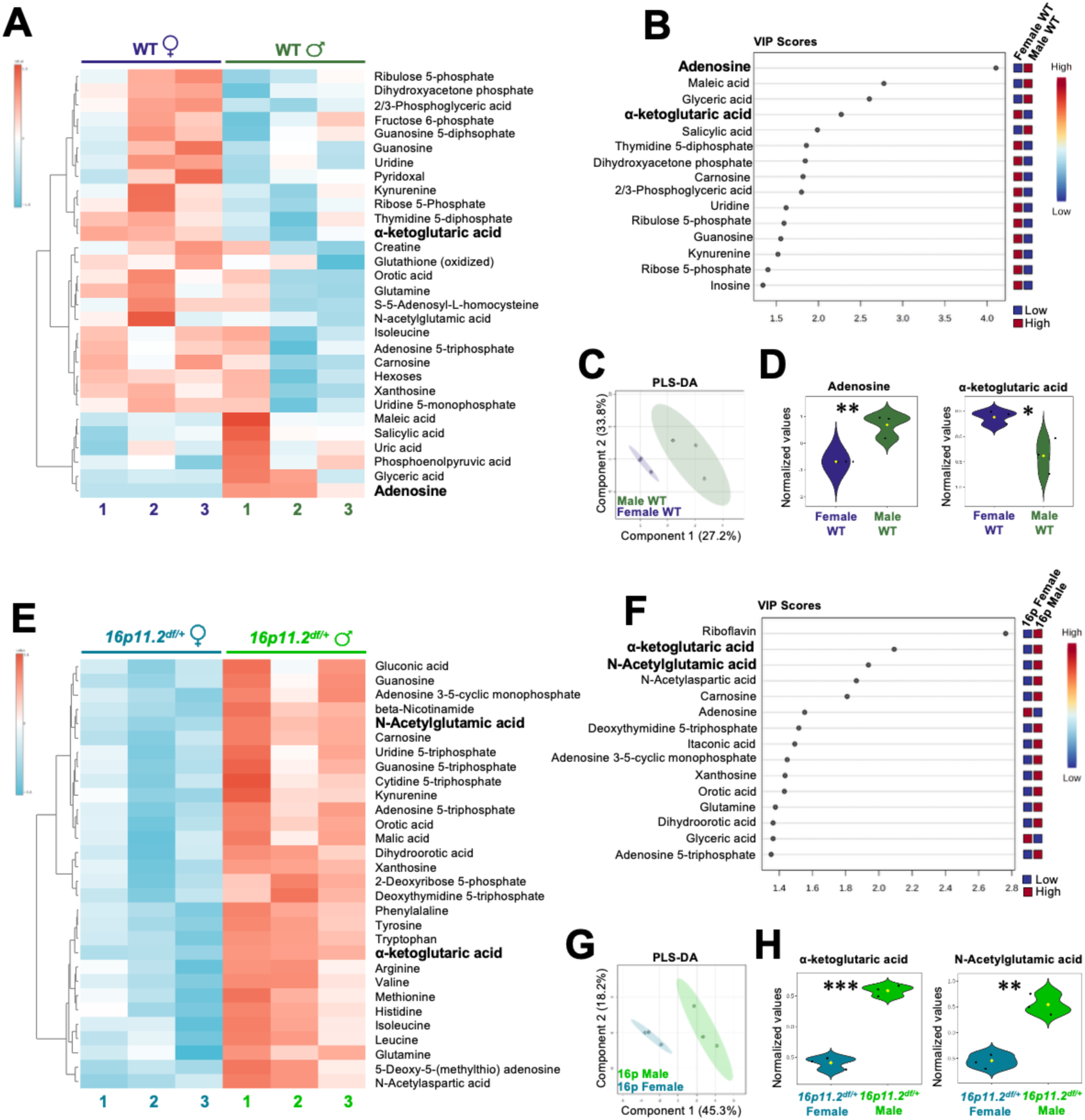
Sex-specific metabolomic profiles conserved in cortical astrocytes from female versus female 16p11.2-deficient WT P8 pups. (A) Heat map (multivariate statistical analysis) displaying the top 30 metabolites significantly altered by genotype from female WT and male WT cortical astrocytes, isolated from 3 mice (n=3) per genotype. (B) Variable importance in projection (VIP) scores, showing the top 15 (score > 1) metabolite drivers contributing to the separation of metabolic profiles identified by Partial Least Squares-Discriminant Analysis (PLS-DA) in female WT and male WT astrocytes. (C) PLS-DA of metabolomics data from female WT (n=3) and male WT (n=3) astrocytes. PLS-DA was obtained with 2 components. The explained variances are shown in parentheses. (D) Details of most significantly altered metabolites from female WT (n=3) and male WT (n=3) astrocytes. Identified with negative log10-transformed p-values from the t-test plotted against the log2 fold change normalized by Pareto scaling. *Left,* Violin plot of adenosine **(p=0.005) (yellow data point represents mean, black data point represents individual mouse). *Right,* Violin plot of alpha-ketoglutaric acid *(p=0.022) (yellow data point represents mean, black data point represents individual mouse). (E) Heat map (multivariate statistical analysis) displaying the top 30 metabolites significantly altered by genotype from female *16p11.2^df/+^* (n=3) and male *16p11.2^df/+^* (n=3) cortical astrocytes. (F) VIP scores, showing the top 15 (score > 1) metabolite drivers contributing to the separation of metabolic profiles identified by PLS-DA in female *16p11.2^df/+^*and male *16p11.2^df/+^* astrocytes. (G) PLS-DA on metabolomics data from female *16p11.2^df/+^* (n=3) and male *16p11.2^df/+^*(n=3) astrocytes. PLS-DA was obtained with 2 components. The explained variances are shown in parentheses. (H) Details of most significantly altered metabolites from female *16p11.2^df/+^* (n=3) and male *16p11.2^df/+^*(n=3) astrocytes. Identified with negative log10-transformed p-values from the t-test plotted against the log2 fold change normalized by Pareto scaling. *Left* Violin plot of alpha-ketoglutaric acid ***(p=0.0001) (yellow dot represents mean, black data points represent individual mice). *Right* Violin plot of N-acetylglutamic acid **(p=0.001) (yellow dot represents mean, black data points represent individual mice).

**Figure S3.**
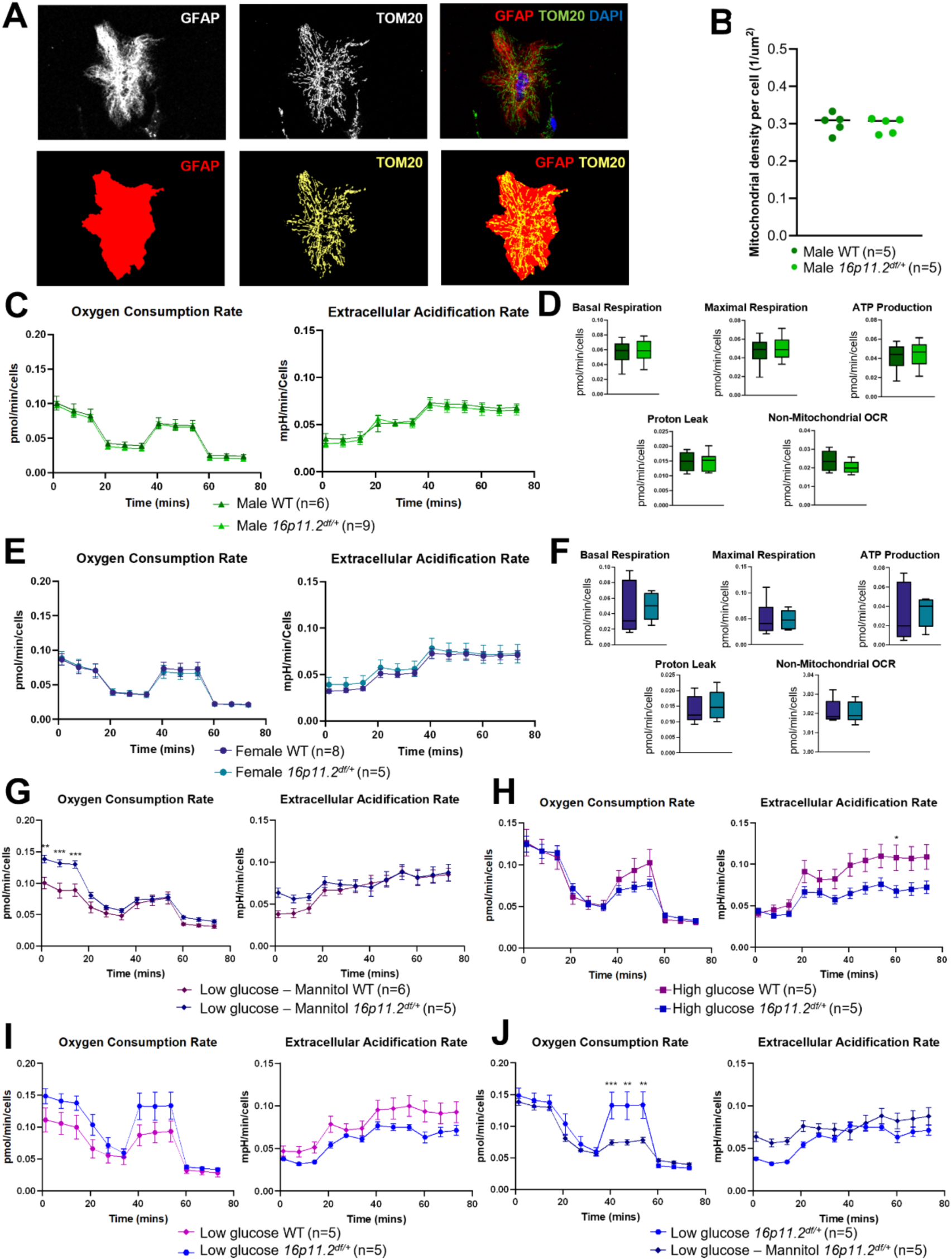
Mitochondrial function in male and female cortical astrocytes at baseline and altered glucose conditions. (A) Representative immunocytochemistry images of primary murine cortical astrocytes stained with GFAP as an astrocytic cytoskeletal marker; TOM20 as a mitochondrial marker. GFAP staining was used to outline the cell surface. TOM20 coverage area was quantified. (B) TOM20 coverage area quantified within astrocyte perimeter as an index of mitochondrial density per cell. No significant difference was identified between *16p11.2^df/+^* (n=5) and WT (n=5) male astrocytes. (C) *Left,* Quantification of oxygen consumption rate (OCR) normalized for cell count. *Right,* Quantification of extracellular acidification rate (ECAR) normalized for cell count. Data are mean ± s.e.m. for male *16p11.2^df/+^* (n=9) and WT (n=6) astrocytes. (D) Quantification of OCR parameters: basal respiration, proton leak, maximal respiration, non-mitochondrial OCR and ATP production were not changed in male *16p11.2^df/+^* astrocytes. Whisker boxes (min to max, center line indicating median), or traces. (E) *Left*, OCR normalized for cell count. *Right,* ECAR normalized for cell count. Data are mean ± s.e.m. for female *16p11.2^df/+^* (n=9) and WT (n=6) astrocytes. (F) Quantification of OCR parameters: basal respiration, proton leak, maximal respiration, non-mitochondrial OCR and ATP production were not changed in *16p11.2^df/+^* female derived astrocytes. Whisker boxes (min to max, center line indicating median), or traces. (G) *Left,* OCR comparison of low glucose (5.5 mM) with mannitol (19.5 mM) normalized for cell count. Data are mean ± s.e.m. combined male and female *16p11.2^df/+^* (n=6) vs WT (n=5) astrocytes. Significant difference was observed in the baseline time measurements at 1.3 min adjusted p value **(p<0.001), at 7.8 min ***(p<0.0001), and 14.3 min ***(p=0.0004) by 2-way repeated measure ANOVA and Sidak’s multiple comparison *post hoc* test. *Right,* ECAR comparison of low glucose (5.5 mM) with mannitol (19.5 mM) normalized for cell count. Data are mean ± s.e.m. combined male and female *16p11.2^df/+^*(n=6) vs WT (n=5) astrocytes. (H) *Left,* OCR comparison of high glucose (25 mM) normalized for cell count. Data are mean ± s.e.m. combined male and female *16p11.2^df/+^* (n=5) vs WT (n=5) astrocytes. *Right,* ECAR comparison of high glucose (25 mM) *16p11.2^df/+^*(n=5) vs WT (n=5) littermates. Significant difference was observed in the maximal respiration measurements at 60.3 mins adjusted p value *(p=0.04) by 2-way repeated measure ANOVA and Sidak’s multiple comparison *post hoc* test. (I) *Left,* OCR comparison of low glucose (5.5 mM) normalized for cell count. Data are mean ± s.e.m. combined male and female *16p11.2^df/+^* (n=5) vs WT (n=5) astrocytes. No significant difference was observed. *Right,* ECAR comparison of low glucose (5.5 mM) *16p11.2^df/+^* (n=5) vs WT (n=5) littermates. No significant difference was observed either. (J) *Left,* OCR comparison of low glucose (5.5 mM) and low glucose (5.5 mM) with mannitol (19.5 mM) normalized for cell count. Data are mean ± s.e.m. combined male and female *16p11.2^df/+^* (n=6) vs WT (n=5) astrocytes. Difference was observed during maximal respiration at 40.6 min ***(p<0.001), at 47.1 min **(p=0.001), and 53.7 min **(p=0.002) by 2-way repeated measure ANOVA and Sidak’s multiple comparison *post hoc* test. *Right,* ECAR comparison of low glucose (5.5 mM) with mannitol (19.5 mM) normalized for cell count. Data are mean ± s.e.m. combined male and female *16p11.2^df/+^*(n=6) vs WT (n=5) astrocytes. No significant difference was observed, by 2-way repeated measure ANOVA and Sidak’s multiple comparison *post hoc* test.

**Figure S4.**
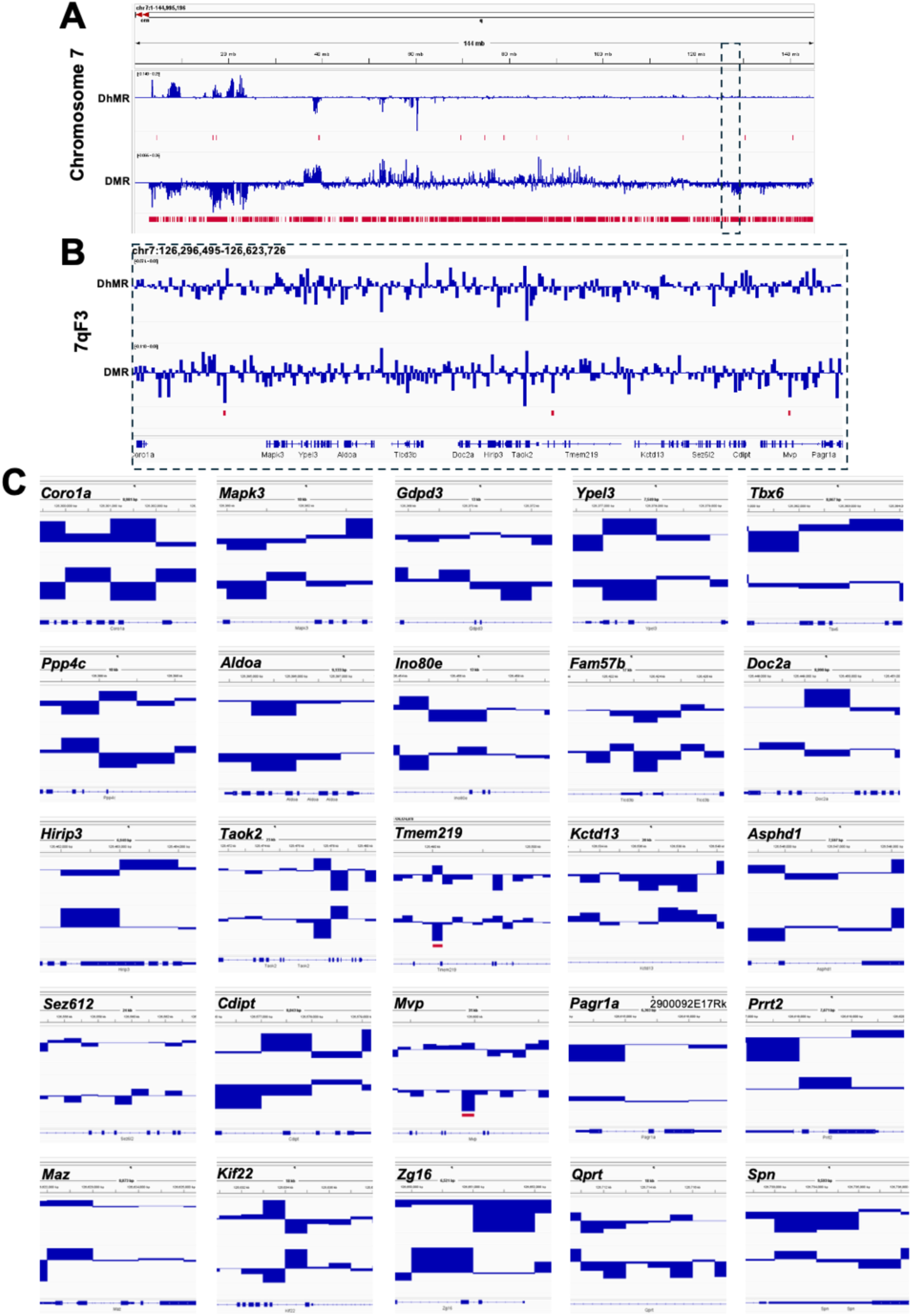
Little to no methylation changes within the 16p11.2 (7qF3) locus. (A) Integrative Genomics Viewer (IGV) snapshot of chromosome 7 hydroxymethylation and methylation differences, , between male *16p11.2^df/+^* (n=4) and WT (n=4) astrocyte DNA. Hydroxymethylation and methylation read differences represented in blue tracks, with Differentially Hydroxymethylated Regions (DhMRs) and Differentially Methylated Regions (DMRs) meeting multiple-comparison-corrected p-value (false discovery rate, FDR of 0.1) represented in red tracks. Bottom track is mouse reference genome (GRCm39/mm39). (B) Detailed IGV representation of genomic region in to the 16p11.2 (7qF3) locus (chr7:126,296,495-126,623,726). (C) IGV snapshots of genes associated with the 7qF3 locus, including *Coro1a* (chr7:126,296,946-126,305,925), *Mapk3* (chr7:126,356,798-126,366,988), *Gdpd3* (chr7:126,363,586-126,376,817), *Ypel3* (chr7:126,374,135-126,381,682), *Tbx6* (chr7:126,378,655-126,386,720), *Ppp4c* (chr7:126,383,038-126,393,719), *Aldoa* (chr7:126,392,406-126,401,537), *Ino80e* (chr7:126,449,605-126,463,544), *Fam57b* (chr7:126,414,057-126,431,391), *Doc2a* (chr7:126,444,889-126,453,877), *Hirip3* (chr7:126,459,611-126,466,549), *Taok2* (chr7:126,462,849-126,486,139), *Tmem219* (chr7:126,483,392-126,524,078), *Kctd13* (chr7:126,526,051-126,546,781), *Asphd1* (chr7:126,543,159-126,550,754), *Sez6l2* (chr7:126,547,707-126,571,778), *Cdipt* (chr7:126,573,630-126,581,671), *Mvp* (chr7:126,584,040-126,615,729), *Pagra1a* (chr7:126,612,223-126,618,524), *Prrt2* (chr7:126,614,714-126,622,383), *Maz* (chr7:126,619,306-126,628,177), *Kif22* (chr7:126,624,901-126,643,579), *Zg16* (chr7:126,647,328-126,653,847), *Qprt* (chr7:126,704,942-126,723,201), and *Spn* (chr7:126,729,404-126,738,995).

**Figure S5.**
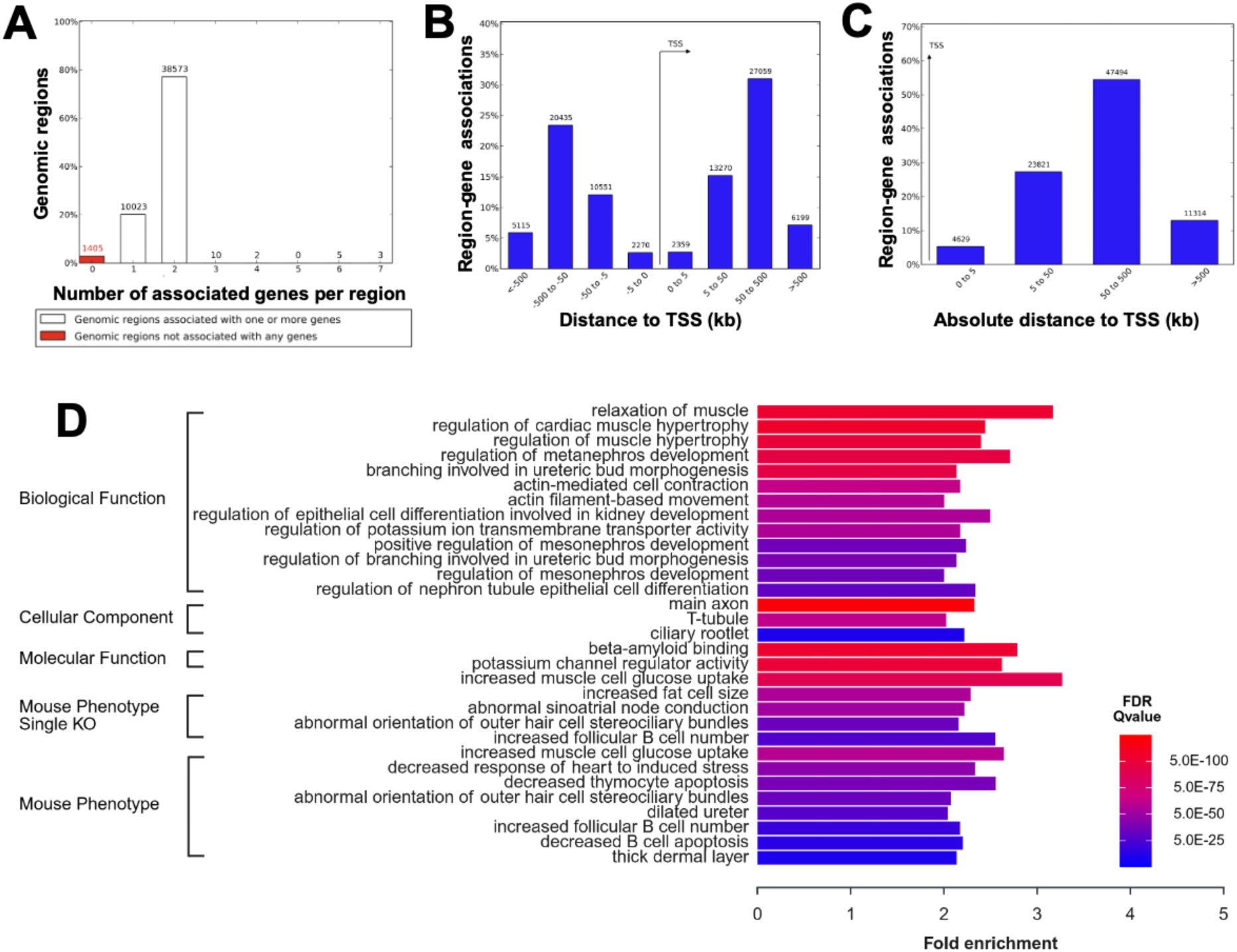
Genomics Regions Enrichment of Annotations Tool (GREAT) analysis for Gene Ontology. (A) Percentage of associated genes by region. (B) Percentage of gene-region associations by orientation and distance to transcript start site (TSS) in kb of the DMR input regions. (C) Percentage of gene-region associations to absolute distance to TSS of the DMR input regions in kb.(D) The GREAT analysis identified gene ontology terms for 50,021 identified differential methyl regions with a statistical significance threshold of FDR 0.1. Test set of 50,021 genomic regions picked 13,838 (65%) of all 21,395 genes. Gene ontology terms were tested using an annotation count range of [1, Inf].

**Figure S6.**
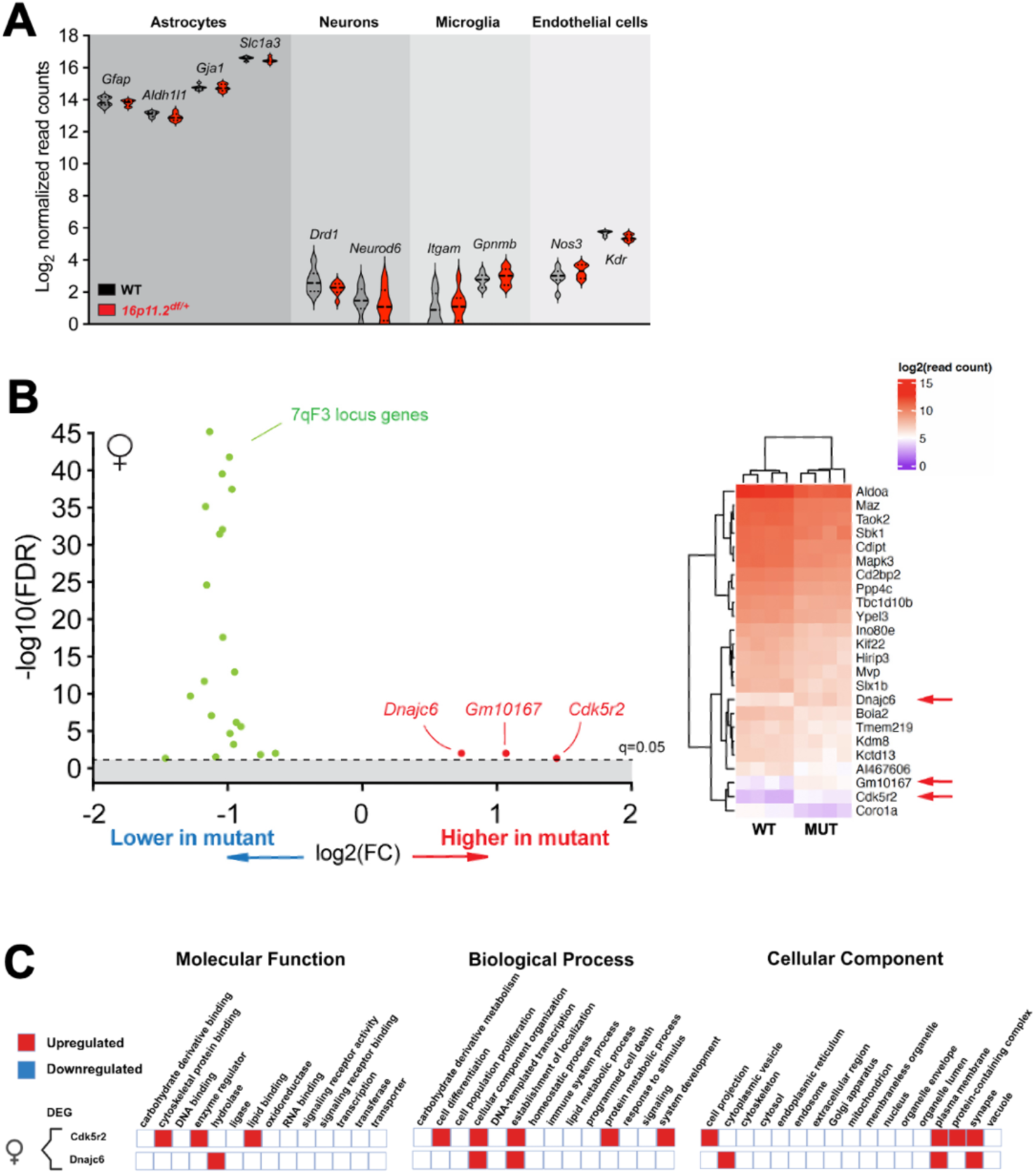
Differential gene expression in female *16p11.2^df/+^* P8 astrocytes. (A) Quality control for acute RNA isolation from cortical extracts via assessment of astroglial gene enrichment. High enrichment of astrocyte-specific genes including *Gfap*, *Aldh1l1, Gja1 or Slc1a3* was confirmed. Low contamination by transcripts from neurons (*Drd1, Neurod6*), microglia (*Itgam, Gpnmb*) or endothelial cells (*Nos3, Kdr*) was noted. Violin plots are from Log2 of normalized read counts. (B) *Left,* Volcano plot of differentially expressed genes (DEGs) identified with false discovery rate (FDR)-adjusted q-value<0.05, sequenced from female *16p11.2^df/+^* (n=4) and WT (n=4) astrocytes (upregulated genes in red, downregulated in blue, 7qF3 locus genes in green). *Right,* Heatmap (multivariate statistical analysis) of DEGs detected in female *16p11.2^df/+^* (n=4) and WT (n=4) astrocytes (upregulated genes indicated with red arrows). (C) Gene Ontology terms associated with female DEGs (upregulated genes).

**Figure S7.**
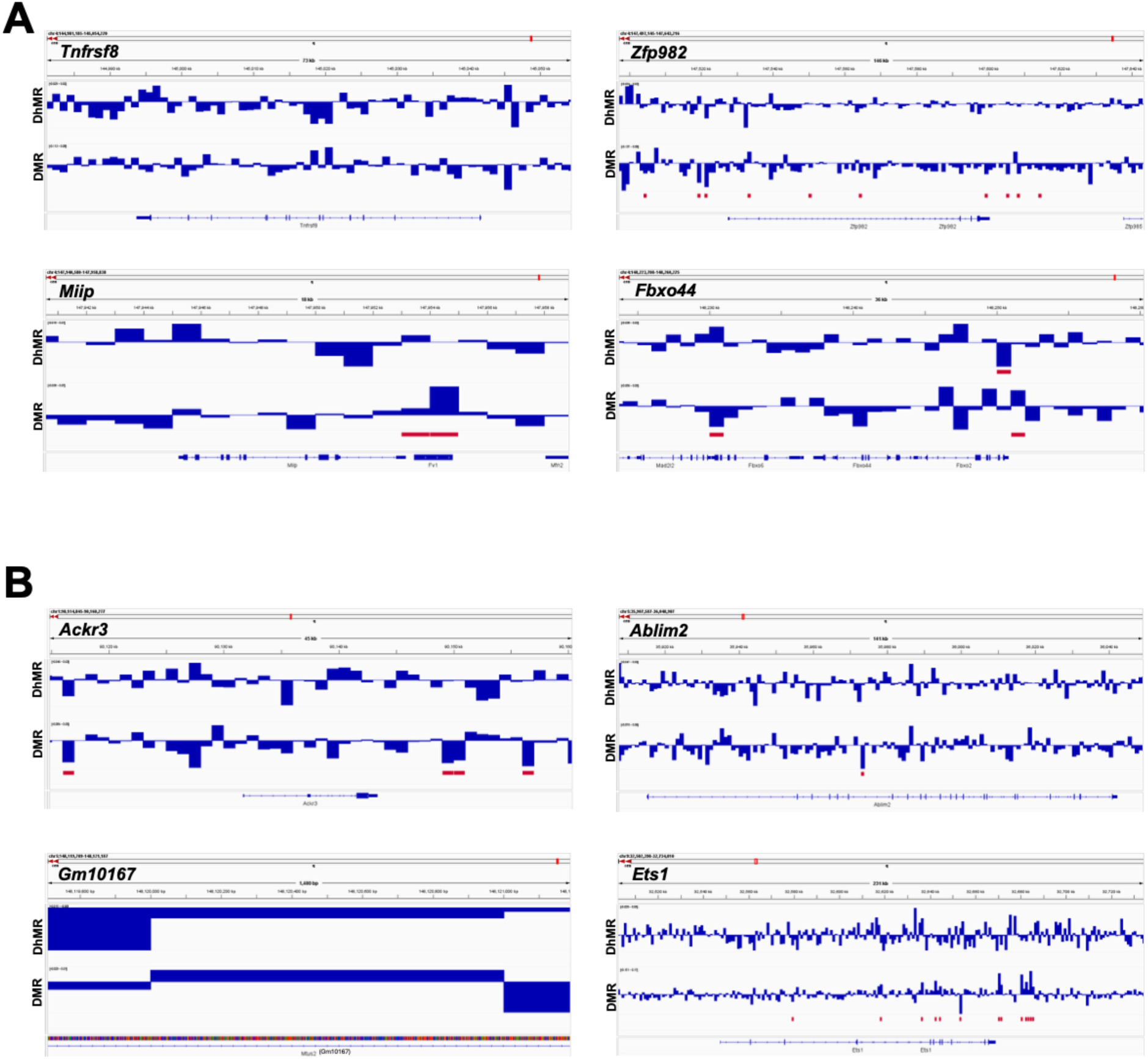
Methylation data for DEGs identified in male *16p11.2^df/+^* astrocytes. (A) Integrative Genomics Viewer (IGV) snapshot of hydroxymethylation and methylation differences between male *16p11.2^df/+^* (n=3) and WT (n=3) astroglial DNA for downregulated DEGs, including *Tnfrsf8* (chr4:144,991,702-145,043,734), *Zfp982* (chr4:147,525,571-147,602,521), *Miip* (chr4:147,943,235-147,955,176) and *Fbxo44* (chr4:148,235,256-148,246,551). (B) IGV snapshot of hydroxymethylation and methylation differences between male *16p11.2^df/+^* (n=3) and WT (n=3) astroglial DNA for upregulated DEGs, including *Ackr3* (chr1:90,129,702-90,145,446), *Ablim2* (chr5:35,913,224-36,044,323), *Gm10167* (Chr5:148119709-148121187), and *Ets1* (chr9:32,545,537-32,671,116).

